# Photocaged FLP Recombinase for Precise Spatio-temporal Control of Gene Expression

**DOI:** 10.1101/2025.05.06.652468

**Authors:** Jose Javier Vazquez Rodriguez, Kieran Baxter, Yunchuan Ma, Sebastian Greiss

**Affiliations:** Institute for Neuroscience and Cardiovascular Research, University of Edinburgh, United Kingdom; Present address: Max Planck Institute for Biological Cybernetics, Tuebingen, Germany

## Abstract

The ability to precisely control gene expression is fundamental to studying biological processes. Using site-specific recombinases such as FLP, gene expression can be controlled, albeit with limited spatiotemporal precision. We develop a photocaged FLP recombinase which can be precisely controlled using light, and we demonstrate its efficacy in *C. elegans.* We use genetic code expansion to incorporate photo-caged amino acids into FLP, replacing critical residues in the active site with their photocaged counterparts. Photocaged FLP displays no detectable background activity, and brief illumination, can be used to activate FLP with near 100% efficiency. We show that photocaged FLP can be activated by light between 365 nm and 435 nm, and that it is not activated by light above 450 nm, making it fully compatible with wavelengths commonly used for imaging and optogenetics. Furthermore, we demonstrate that photocaged FLP can be used to switch on expression of target genes in individual cells within the animal using a standard 405 nm microscope mounted laser to deliver the activating light. Activation by laser requires illumination times of less than 10 ms per cell. Thus, we have developed a straightforward and efficient tool to precisely control gene expression in the multicellular organism *C. elegans*.

## INTRODUCTION

Methods to control gene expression are fundamental to studying biological processes in living systems. Such methods include the use of tissue-specific promoters or of extraneous transcription factors such as Gal4 to drive the expression of target genes^1^. Another widely used approach involves site specific DNA recombinases (SSR) such as Cre or FLP, which can remodel DNA to switch the expression of target genes on or off^2^. However, spatial and temporal targeting of SSR systems depends on the availability of promoters to drive SSR expression. This is limiting as often promoters with desirable spatial and temporal expression profiles for an experiment do not exist. Several strategies have been employed to overcome this limitation.

Temporal control can be achieved through chemically inducible systems, for example using tamoxifen^3^, or by employing heat-shock inducible promoters^4^. Spatial control can be improved through the use of overlapping promoter pairs to express individual components of the system^5^.

More recently, approaches to optically control the activity of SSRs have been developed. Optical activation is non-invasive and has the potential to offer unmatched spatial and temporal control. Most approaches for imparting optical responsiveness to SSRs involve split proteins that are reassembled using photodimerizers such as LOV domains^6^. While this approach has been applied for Cre^7^ and FLP^6^ recombinases, it is often accompanied by background activity in the ‘off’ state, and low efficiency in the ‘on’ state. Additionally, its practical utility is hindered by the requirement for prolonged light exposure to induce recombinase reconstitution, and the use of wavelengths in the visible spectrum which overlap with wavelengths often used for imaging and optogenetic applications.

An alternative approach for optically controlling SSR activity involves the incorporation of photocaged amino acids into the enzyme’s active site in place of essential catalytic residues using genetic code expansion (GCE) technology^8^. Here, a photolabile caging group attached to the ncAA side-chain blocks activity of the enzyme, until brief illumination removes the caging group and restores protein activity. This approach has been successful for Cre, which has been photocaged using photocaged variants of tyrosine to replace the active site Tyr324^9,10^, and photocaged variants of lysine to replace the active site Lys201^10–12^. The use of photocaged amino acids has several advantages: the photocaging groups are stable at wavelengths commonly used for imaging and other optogenetic applications; photocaged Cre (PC-Cre) shows no background activity prior to illumination; activation can be achieved with sub-second illumination time; and the light can be precisely targeted to individual cells, thereby allowing the control of gene expression with single-cell precision^12^. PC-Cre has been successfully applied in single-celled systems^9,10^ as well as in animals such as zebrafish^11^ and *C. elegans*^12^.

Several SSRs besides Cre exist and have been employed to control gene expression in cells and multicellular organisms^2^. The most widely used among these is flippase (FLP), originally derived from the 2 μ plasmid of *S. cerevisiae*^13^. Compared to Cre, FLP is more active at the lower body temperatures of fish, worms, and flies^14^. At the same time, versions of FLP with improved activity and stability at 37 °C for use in mammalian systems have been developed^15^. Furthermore, while Cre can be toxic to cells^16^, no such deleterious effects have been reported for FLP^17^.

Here, we develop photocaged FLP (PC-FLP) by replacing specific tyrosine, lysine, and cysteine residues in its active site with photocaged variants. We find that FLP can be photocaged with no background activity and that it can be activated with up to 100% efficiency upon illumination. We also find that in *C. elegans,* PC-FLP is more efficient than PC-Cre. Furthermore, we have identified PC-FLP variants with improved activation efficiency at longer wavelengths and demonstrated efficient activation using a standard 405 nm microscope-mounted laser and illumination times of less than 10 ms per cell, enabling the control of gene expression with single cell precision in *C. elegans*.

## METHODS

### C. elegans strains

Strains were maintained under standard conditions unless otherwise indicated^18,19^. Transgenic strains were generated by biolistic bombardment using hygromycin B as a selection marker^12,20,21^. Strains were maintained on Nematode growth medium (NGM) supplemented with hygromycin at a concentration of 0.3 mg/ml (Formedium).

### Plasmid construction

Expression plasmids transformed into *C. elegans* were assembled from entry plasmids and destination vectors using the Gateway LR Clonase II Plus Enzyme Mix (Thermo Fisher Scientific). DNA fragments and oligonucleotides used in the generation of entry plasmids were custom synthesised by IDT. Point mutations were introduced using PCR mutagenesis. Description and construction of expression plasmids is detailed in Supplementary Table 2. Entry plasmids used in the LR reactions are described in Supplementary table 3. The sequences for the genes described in this work are given in Supplementary Note 1.

### ncAA feeding

PCK^22^ was custom synthesised by ChiroBlock GmbH, Germany. HCK^23^ was custom synthesised by Inochem, United Kingdom. NPY^24^ was custom synthesised by NewChem Technologies, UK. MNPY^24^ was custom synthesised by NewChem Technologies, UK. ONBY^25^ was purchased from Fluorochem (product code F447104). PCC2^26^ was custom synthesised by ChiroBlock GmbH, Germany. For commercial availability of ncAAs used in this study see Supplementary Note 2.

NGM plates containing ncAA (ncAA-NGM) were prepared by dissolving the ncAA in a small volume of HCl or NaOH and then adding it to the molten NGM. The HCl or NaOH were then neutralised by equimolar NaOH or HCl respectively. PCK was dissolved in 0.2 M HCl, HCK was dissolved in 0.1 M NaOH, NPY was dissolved in 1 M NaOH, MNPY was dissolved in 0.1 M NaOH, ONBY was dissolved in 1 M NaOH, PCC2 was dissolved in 0.5 M HCl.

Animals were age-synchronised by settling and added to ncAA-NGM plates as L1 larvae. Freeze-dried OP50 (LabTIE) reconstituted according to the manufacturer’s instructions was added for food. Animals were grown on the ncAA-NGM plates for 24 hours unless otherwise stated.

### Photocaged Recombinase activation

For global activation, worms were washed off NGM plates with M9 supplemented with 0.001% Triton-X100. To test activation, animals were illuminated in a CL-1000L crosslinker at 365 nm (Analytik Jena) for 5 minutes. To test activation at defined longer wavelengths, a pE-4000 light source (CoolLED) was used.

Worms were illuminated with the “365 nm”, “405 nm”, “435 nm”, “460 nm”, or “470 nm” LEDs at 100% power for 5 minutes. These LEDs produce light in the following ranges: “365 nm” (350 nm to 415 nm), "405 nm" (370 nm to 443 nm), "435 nm" (403 nm to 470 nm), "460 nm" (427 to 510 nm) and "470 nm" (433 nm to 522 nm) (see also **Supplementary Fig. 9b**), and are rated at 30 mW, 188 mW, 170 mW, 241 mW and 167 mW respectively.

For laser activation at 405 nm, a Zeiss LSM800 laser scanning confocal microscope was used. Worms were anaesthetized with 50 mM NaN3 and mounted on 2.5% agar pads on microscope slides. For whole-head illumination, the worms were located with the 20x objective, and the bleaching function was run for 10 iterations with a 2.31 µs dwell time at 2 µm Z-stack intervals across the entire width of the worm. For single-cell illumination, worms were located with the 20x objective and a 6x area crop was applied. The bleaching function was run for 20 iterations with a 2.31 µs dwell time at a single Z-plane within a ‘Region of Interest’ that was drawn around the cell body of the target neuron. Based on the known pixel size at 20x magnification (0.7 µm), the dwell time (2.31 µs), bleaching cycles used (20), the total illumination time can be calculated to be in the range of 1ms to 6ms per cell.

The laser power levels at 50%, 10%, and 1% were measured to be 632.93±17.91 µW, 127.76±1.24 µW, and 11.34±0.17 µW respectively using a PM100D (Thorlabs) under the same conditions as used for illuminating the worms. The ± values represent the SEM.

### Imaging and analysis

All imaging was carried out on a Zeiss Axio Imager M2. Worms were anaesthetized with 50 mM NaN_3_ and mounted on 2.5% agar pads on microscope slides. All image analysis was carried out on Fiji version 2.14.0. To score activation efficiency, z-stacks were acquired using a 10x objective and from each stack a single 2D image was generated by sum intensity projection. The fluorescent mKate2 marker within worms was first automatically identified using Li thresholding. This thresholding was used to create a mask that was applied onto the fluorescent Citrine2 reporter channel. Within this mask, the number of pixels that exceeded the signal threshold were counted as signal. The signal threshold for the Citrine2 channel was determined using strain SGR197, which expresses all genetic components except for the Citrine2 reporter construct (Supplementary Table 1). The same signal threshold was used for all images analysed. Activation efficiency was calculated as the ratio of the number of signal pixels to mask pixels, expressed as a percentage. All image analysis steps involved in this analysis were carried out automatically using a Fiji script (Supplementary Note 3). For all conditions at least two independent transgenic strains were analysed and experiments were performed in triplicate.

The automated analysis approach was validated by comparison to a direct cell count. For the direct cell count, cells expressing a respective recombinase were first identified through the mKate2 marker and presence of a nucleus as determined by DIC. Activation efficiency after uncaging was then determined by dividing the number of Citrine2 reporter expressing cells by the total number of recombinase expressing cells. The same images of animals used for direct cell counting were put through the automated analysis. We found no statistically significant difference between the activating efficiency determined by direct cell counting and by the automated analysis (**Supplementary Fig. 1**).

Statistical analyses were carried out using GraphPad Prism version 10. Representative images shown in the figures, except Figure 3 d, were taken using an Apotome 3 module mounted on the Zeiss Axio Imager M2. Images for quantification were taken without the Apotome 3 module.

### Heat shock induced reporter expression

Animals were grown on 1mM NPY for 24h and illuminated in a CL-1000L crosslinker at 365 nm (Analytik Jena) for 5 minutes. 12h after illumination animals were heat shocked at 34 °C for 4h, then returned to 20 °C and imaged 10h after start of the heat shock.

### Behavioural assays

Worms were grown on ncAA-NGM plates for 48 hours. They were then illuminated with 365 nm light for 5 minutes or kept in the dark, after which they were transferred to NGM plates seeded with 20 µl of 0.35 OD *E. coli* OP50 supplemented with ATR to a concentration of 100 µM. Worms were grown in the presence of ATR for 24 hours. Behavioural assays were then carried out on a WormLab setup (MBF Bioscience). Videos of animals illuminated using the integrated 617 nm LED at 5 W/cm² for 1 second were recorded and animals were scored manually as responding or not responding to the optogenetic stimulus. Animals that accelerated forwards or reversed immediately following the stimulus were counted as responders. Animals that did not accelerate or reverse in response to the stimulus were scored as non-responders.

### Recording of ncAA spectra

To measure the UV-Vis spectra of the photocaged ncAAs, they were firstly dissolved in HCl or NaOH as described above. The dissolved ncAA was then added to 25mM potassium phosphate buffer (pH6) to a final concentration of 0.5 mM. UV-Vis spectra were recorded on a Nanodrop 2000 (Thermo Fisher Scientific).

## RESULTS

### Strategy for photocaging FLP recombinase

To photocage FLP, we used genetic code expansion, which enables the cellular translational machinery to utilise, and incorporate into proteins, non-canonical amino acids (ncAAs) with properties not found in nature. This is achieved by expressing an orthogonal aminoacyl-tRNA-synthetase (aaRS)/tRNA pair in the cell. The aaRS specifically recognises the ncAA and attaches it to its cognate tRNA, which in turn delivers it to the ribosome where it is incorporated into the nascent polypeptide chain in response to a designated "unassigned" codon, most commonly a UAG (amber) stop codon (**Fig. 1a**). Here we utilise the pyrrolysyl-tRNA-synthetase (PylRS)/tRNA(Pyl) pair from the archaeon *Methanosarcina mazei*, which is highly efficient in eukaryotic cells and the most commonly used pair in in multicellular organisms^27^.

**Fig. 1.**
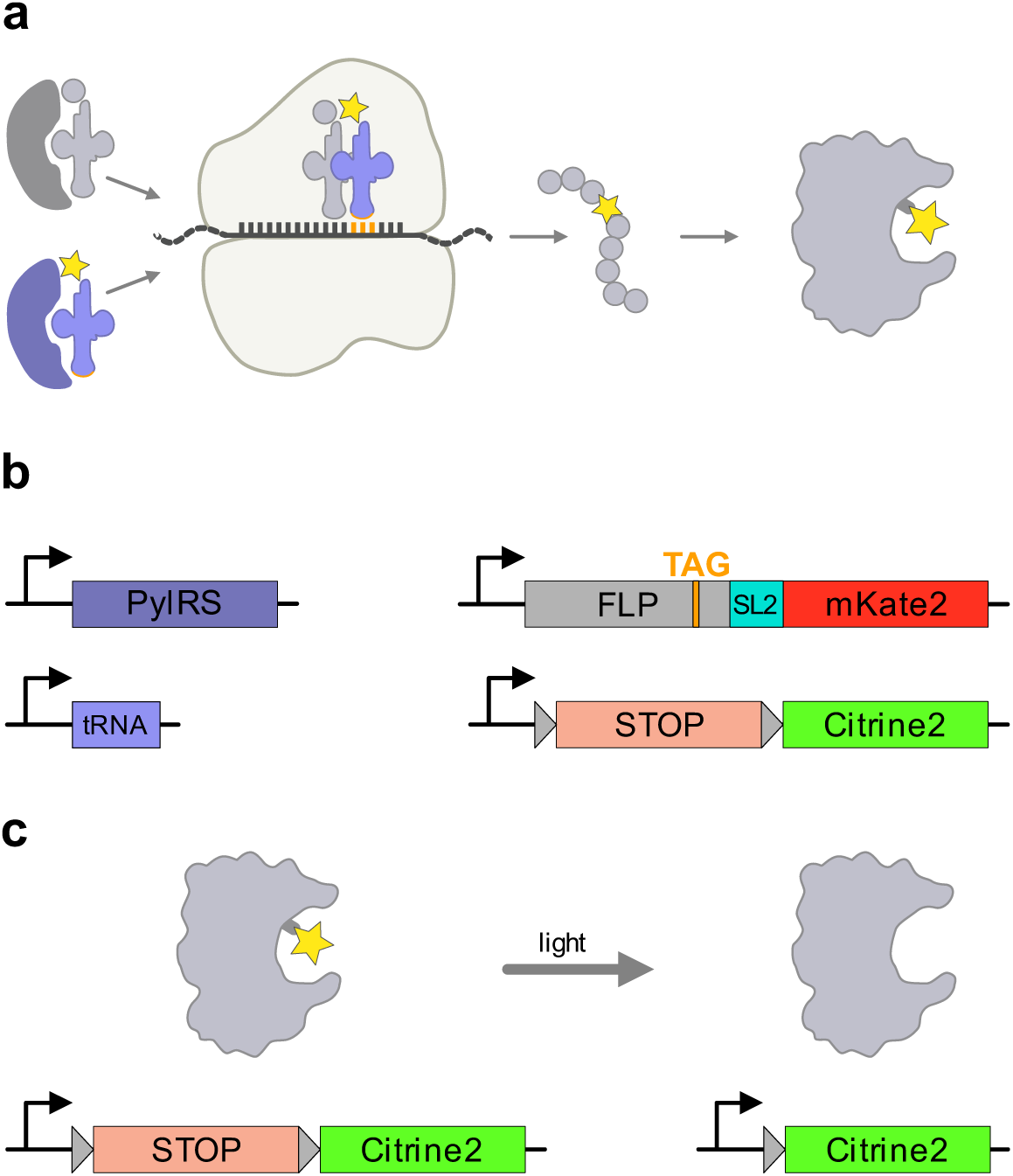
Photocaging the activity of FLP recombinase using genetic code expansion. **a** The orthogonal PylRS aminoacylates its cognate orthogonal tRNA (blue pair) with the ncAA (yellow star). The amino-acylated orthogonal tRNA then directs the incorporation of the ncAA into the growing polypeptide chain in response to the amber stop codon. This results in the incorporation of the photocaged ncAA into FLP to produce PC-FLP. **b** Constructs transformed into *C. elegans* for expression and visualisation of PC-FLP. The PylRS and tRNA serve to direct the incorporation of the various photocaged ncAAs into the FLP gene at the amber stop codon. FLP is expressed in an operon with mKate2 to visualise its expression. The reporter is made up of a Citrine2 gene whose expression is prevented by an upstream transcriptional stop cassette that is flanked by FRT sites. **c** PC-FLP, which is photocaged by the incorporation of a photocaged ncAA into its active site is unable to excise the transcriptional stop cassette inserted between the Citrine2 reporter gene and its promoter. Following illumination, the photocaging group within PC-FLP is removed, producing active FLP, which is able to excise the transcriptional terminator and allow for the production of Citrine2.

To qualify as a target for photocaging, a residue in an enzyme needs to fulfil two criteria: installing the photocaged ncAA must abolish enzymatic activity, and removal of the caging group must restore activity. To select residues for photocaging FLP that fulfil these criteria, we first compared its structure to that of Cre, which has previously been successfully photocaged.

Although Cre and FLP share only 13% sequence identity and assemble their active sites differently, both belong to the tyrosine recombinase family and therefore contain the conserved catalytic pentad^2^, which includes the residues K201 and Y324 at which Cre has previously been photocaged. These residues are conserved in FLP as K223 and Y343^2,28^. Therefore, we investigated K223 and Y343 as targets for installing optical control in FLP by replacing them with their photocaged counterparts (**Fig. 2b**, **Supplementary Fig. 2**).

**Fig. 2.**
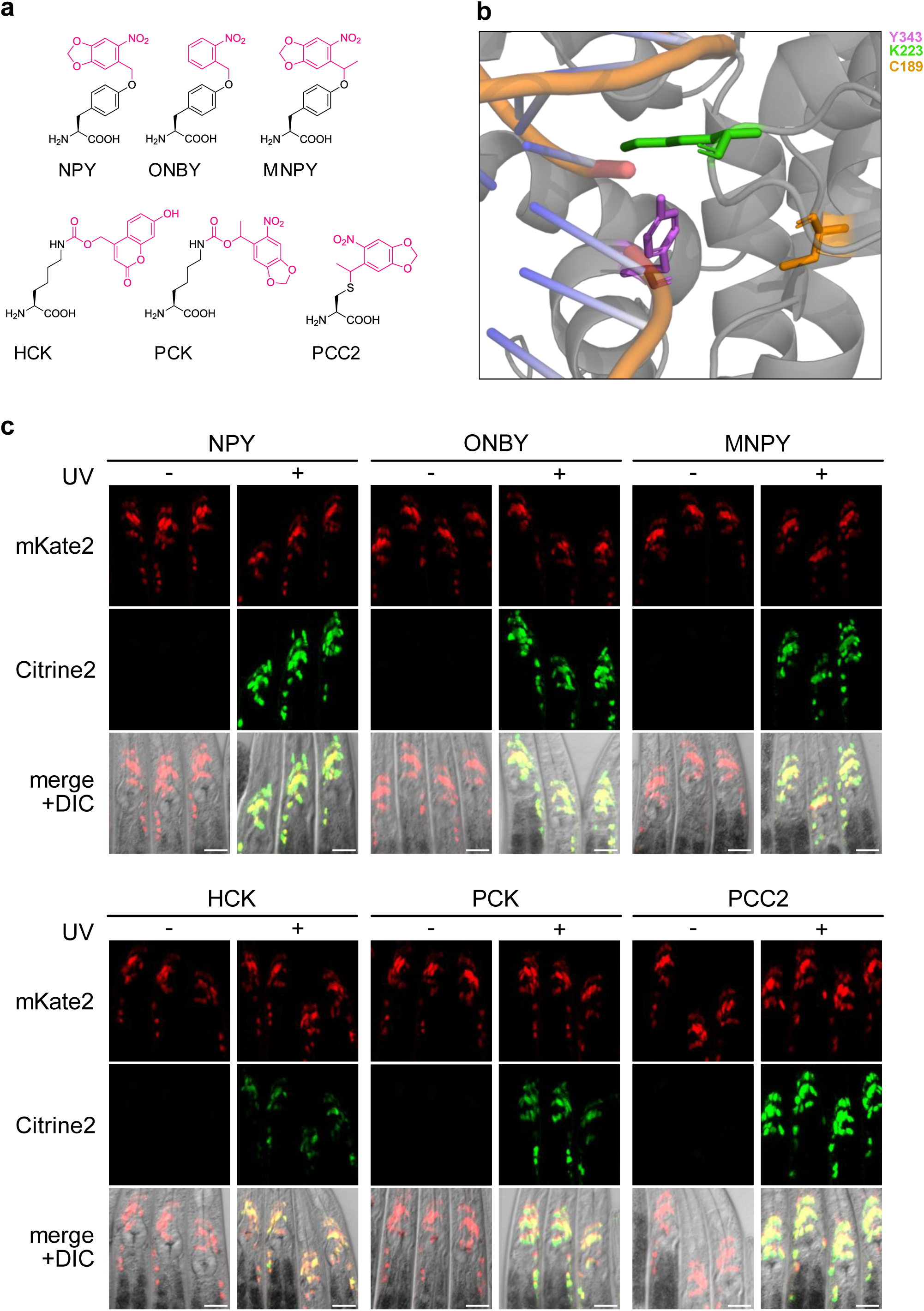
PC-FLP system. **a** Chemical structures of photocaged ncAAs used in this paper. The photocaging groups are highlighted in pink. **b** Active site of FLP recombinase (PDB: 1FLO^35^) with active site residues Y343, K223, and C189 highlighted in purple, green, and orange respectively. **c** Fluorescence microscopy images of *C. elegans* head regions expressing the PC-FLP system using the different photocaged variants of tyrosine, lysine, and cysteine. Worms were grown in the presence of the respective ncAA at a concentration of 1 mM for 24 hours and then illuminated for 5 minutes with 365 nm ("+" condition) or kept in the dark ("-" condition). Scale bars represent 20 µm.

In addition to the conserved catalytic tyrosine and lysine residues that are shared between FLP and Cre, we also identified C189 that is present in FLP, but not in Cre, as a photocaging target. FLP belongs to the family of yeast-derived tyrosine recombinases, which share between them greater sequence conservation than FLP does with Cre and contain several invariant residues. C189 is one of these invariant residues, located within the active site of FLP, and which if mutated to arginine or tryptophan results in a complete loss of function^29^. We have previously shown that the photocaged cysteine PCC2 can be efficiently incorporated in *C. elegans* using the evolved PylRS variant PCC2RS^26,30^. Thus, we chose C189 as another promising candidate residue for photocaging the activity of FLP (**Fig. 2b**, **Supplementary Fig. 2**).

To express and assay photocaged FLP (PC-FLP) variants, we generated at least two independently derived transgenic *C. elegans* strains for each of the FLP constructs containing an amber stop codon at the position that corresponds to the ncAA incorporation site (**Fig. 1b**, **Supplementary Table 1**). Along with the FLP construct, we co-transformed the genetic code expansion machinery components, namely a PylRS variants evolved to recognise the respective photocaged ncAA and optimised for use in eukaryotic cells by the addition of a N-terminal nuclear export signal^12,31^, and tRNA(M15)_CUA_, a previously optimised pyrrolysyl tRNA^12,32^ to decode the amber stop codon present in FLP (**Fig. 1b**). Expression of the PylRS and FLP variants was driven by the promoter for the glutamate receptor *glr-1* that is active in a subset of *C. elegans* head neurons^12^. To visualise presence of the expression construct in our transgenic lines, we co-expressed red fluorescent mKate2 in an artificial operon with FLP. The FLP gene was modified along the lines of our previously published optimised PC-Cre by adding a C-terminal nuclear localisation signal (NLS) derived from *C. elegans egl-13*^12^. The C-terminal NLS facilitates nuclear translocation of full length FLP. As a reporter to visualise FLP activity we co-transformed a construct consisting of a Citrine2 gene separated from its promoter by a transcriptional terminator flanked by two FRT sites. FLP activity would thus result in excision of the transcriptional terminator and expression of the fluorescent Citrine2 (**Fig. 1c**). To drive expression of the Citrine2 reporter we used the promoter of *C. elegans maco-1*, which is active pan-neuronally^12^.

Using this strategy, namely transgenic animals that co-express FLP constructs containing an amber stop codon in the selected position, the genetic code expansion machinery components, and the reporter system for visualising recombinase activity, we assessed the photocaging efficiency in the three selected FLP residues.

### Photocaging FLP at Tyrosine 343

To photocage FLP at the catalytic Y343, we tested three different photocaged tyrosine variants. The photocaged tyrosine ortho-nitro-benzyl tyrosine (ONBY), which was previously used to photocage Cre^10^, and the photocaged tyrosine variants nitropiperonyl-tyrosine (NPY) and methyl-nitropiperonyl-tyrosine (MNPY) (**Fig. 2a**). NPY and MNPY show improved uncaging characteristics compared to ONBY^24^, but have thus far not been used for photocaging either Cre or FLP. We have previously shown that these photocaged tyrosine variants can be efficiently incorporated in *C. elegans* using the evolved PylRS variant NPYRS^24,33^. We therefore evaluated their suitability for photocaging the activity of FLP recombinase.

We generated transgenic *C. elegans* strains by co-expressing NPYRS, tRNA(M15)_CUA_, the FLP construct containing an amber stop codon at position Y343, and the Citrine2 reporter to visualise activity of the recombinase.

To assay activation of PC-FLP using the photocaged tyrosine variants, transgenic animals were grown from the L1 larval stage for 24 hours on Nematode Growth Medium (NGM) agar plates supplemented with 1 mM of either NPY, ONBY, or MNPY, followed by illumination with 365 nm light. 24 hours after illumination we observed strong Citrine2 fluorescence in animals grown in the presence of any of the three photocaged tyrosine variants (**Fig. 2c**), indicating that the function of FLP was restored after illumination. We observed activation efficiencies following illumination of 99.1% 99.5% and 98.5% for PC-FLP photocaged with NPY, ONBY, or MNPY respectively (**Fig. 3a**). In contrast, we saw no Citrine2 expression in animals grown on any of the photocaged tyrosine variants but not illuminated, indicating that all the photocaged tyrosine variants completely abrogated FLP activity when inserted in place of Y343. Likewise, we saw no Citrine2 expression in animals grown without ncAA (**Supplementary Fig. 3**).

**Fig. 3.**
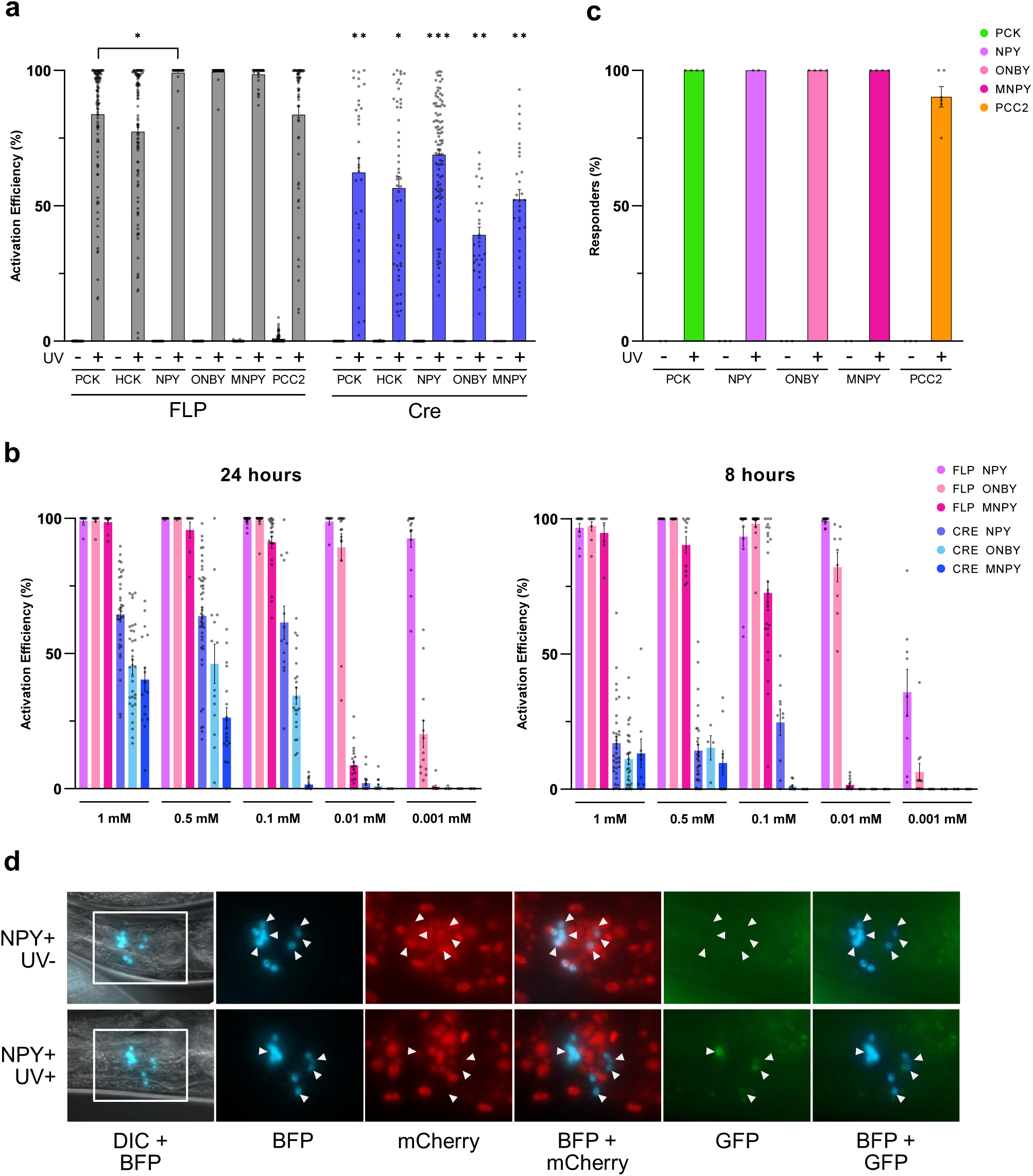
Characterisation of PC-FLP and PC-Cre variants. **a** Activation efficiency in animals expressing PC-FLP and PC-Cre systems using photocaged variants of tyrosine, lysine, and cysteine. Animals were grown in the presence of the respective ncAA at a concentration of 1 mM for 24 hours and then illuminated with 365 nm light ("+" condition) or kept in the dark ("-" condition). Images were taken for quantification 24 hours after illumination. Dots represent individual animals assayed. Error bars represent the SEM. Significance values between conditions were determined by nested t-test carried out to account for the different transgenic strains and repeats. Quantifications from each transgenic strain and repeat are given in Supplementary Fig. 5. The total number of animals for the conditions compared are as follows: UV PC-FLP(K223PCK) n=108, PC-FLP(K223HCK) n=103, PC-FLP(Y343NPY) n=34, PC-FLP(Y343ONBY) n=37, PC-FLP(Y343MNPY) n=39, PC-FLP(C189PCC2) n=61, PC-Cre(K201PCK) n=33, PC-Cre(K201HCK) n=51, PC-Cre(Y324NPY) n=94, PC-Cre(Y324ONBY) n=29, PC-Cre(Y324MNPY) n=32. **b** Activation efficiency in animals expressing the PC-FLP and PC-Cre systems using the three photocaged tyrosine variants NPY, ONBY, and MNPY at decreasing concentrations. The panel on the left shows activation efficiency for worms exposed to the ncAAs for 24 hours. The panel on the right shows activation efficiency for worms exposed to the ncAAs for 8 hours. Dots represent individual animals assayed. Error bars represent the SEM. **c** PC-FLP was used to activate expression of the optogenetic channel Chrimson. Graphs show the percentage of animals expressing the PC-FLP system using the different photocaged variants of tyrosine, lysine, and cysteine that respond to a 1 second pulse of 5 W/cm^2^ 617 nm light 24h after FLP activation using 365 nm light. Worms were grown in the presence of the respective ncAA at a concentration of 1 mM for 48 hours and then illuminated with 365 nm ("+" condition) or kept in the dark ("-" condition). The worms were then transferred to plates supplemented with ATR and behavioural response assayed 24 hours later. Error bars represent the SEM. **d** Fluorescence microscopy images of worms expressing PC-FLP(Y343NPY) that were exposed to 365 nm light or kept in the dark and had genomic reporter expression from the *hsp-16.41* promoter induced by heat shock. The reporter results in expression of GFP::*his-58* or mCherry::*his-58* in cells in which FLP-mediated recombination occurs, or does not, respectively. Arrows in both sets of images point to cells expressing the reporter. Box in the DIC BFP image represents the area cropped into in the other images.

### Photocaging FLP at Lysine 223

To photocage FLP at the catalytic lysine K223, we tested two different photocaged lysine variants. The photocaged lysine variants 6-nitropiperonyl-L-Lysine (PCK) and hydroxycoumarin-lysine (HCK) have previously been used to photocage Cre activity by us and others^10–12^ (**Fig. 2a**). We have previously demonstrated the efficient incorporation of PCK in *C. elegans* using the evolved PylRS variant PCKRS^12,22^. HCK on the other hand has not previously been established for use in *C. elegans*. We nevertheless wanted to explore its suitability for photocaging FLP activity, since HCK has been reported to display efficient uncaging at longer wavelength^23^.

We first tested whether HCK can be incorporated into proteins in *C. elegans* by constructing strains ubiquitously expressing HCKRS, a PylRS variant evolved for incorporating HCK^23^, together with tRNA(M15)_CUA_ and a fluorescent GFP::mCherry reporter containing an amber stop codon between the GFP and mCherry reading frames (**Supplementary Fig. 4a**)^21^. We found that HCK can indeed be incorporated in *C. elegans* efficiently by HCKRS, comparable in efficiency to PCK incorporation using PCKRS (**Supplementary Fig. 4b**).

Interestingly, when we tested cross-reactivity of PCKRS and HCKRS with the two lysine variants, we found that HCKRS efficiently recognized PCK, resulting in incorporation levels comparable to those observed with HCK (**Supplementary Fig. 4b**). In contrast, although PCKRS was capable of incorporating HCK, its efficiency only reached 50% compared to that achieved with PCK. Notably, in strains expressing HCKRS we observed low but clearly detectable production of full length GFP::mCherry protein even in the absence of either ncAA, indicating non-specific amber stop codon read-through in our system and potential incorporation of a canonical amino acid. By comparison, no such full-length reporter protein was detected in strains expressing PCKRS grown in the absence of ncAA. We therefore explored whether we could improve the efficiency of PCKRS so it can be used for incorporation of both photocaged lysine variants PCK and HCK.

To improve efficiency of PCKRS, we tested mutations recently developed for the *M. mazei* PylRS using directed evolution that significantly increase the activity of the enzyme, but do not affect its ncAA specificity^34^. We introduced these mutations to PCKRS to create PCKRS(13IPYER), and found that PCKRS(13IPYER), while it did not show a significant improvement for incorporation of PCK, doubled incorporation levels for HCK (**Supplementary Fig. 4b**). Importantly, PCKRS(13IPYER) retained its fidelity in the absence of ncAA and did not result in production of full length GFP::mCherry without ncAA. We therefore used PCKRS(13IPYER) for further experiments involving either PCK or HCK.

We then generated transgenic *C. elegans* strains expressing PCKRS(13IPYER) together with tRNA(M15)_CUA_, the FLP construct containing an amber stop codon at position K223, and the fluorescent Citrine2 reporter as described above.

To assay activation of PC-FLP using photocaged lysines, transgenic animals were grown from the L1 larval stage for 24 hours on NGM agar plates containing either 1 mM PCK or HCK followed by illumination with 365 nm light to remove the caging group and thus activate FLP. 24 hours after illumination we observed strong Citrine2 fluorescence in animals grown on either PCK or HCK (**Fig. 2c**). We observed activation efficiencies 83.8% and 77.4% when FLP was photocaged by PCK and HCK respectively using PCKRS(13IPYER) (**Fig. 3a**). In contrast activation efficiencies of 31.4% and 71.2% were observed when the HCKRS was used to incorporate PCK and HCK respectively (**Supplementary Fig. 4c**). We saw no Citrine2 expression in any animals grown on either PCK or HCK but not illuminated, indicating that both PCK and HCK were able to completely abrogate FLP activity when inserted in place of K223. Likewise, we saw no Citrine2 expression in animals grown without ncAA (**Supplementary Fig. 3**).

### Photocaging FLP at Cysteine 189

To photocage FLP at C189 we generated transgenic *C. elegans* strains expressing PCC2RS, a PylRS variant evolved to recognise the photocaged cysteine PCC2^26^ (**Fig. 2a**), tRNA(M15)_CUA_, FLP containing an amber stop codon at position C189, and the Citrine2 reporter to visualise activity of the recombinase.

To assay activity of PC-FLP caged using PCC2, transgenic animals were grown from the L1 larval stage for 24 hours on NGM agar plates containing 1 mM PCC2 followed by illumination with 365 nm light to remove the caging group and thus activate FLP. 24 hours after illumination we observed strong Citrine2 fluorescence in animals grown on PCC2 (**Fig. 2c**), corresponding to an activation efficiency of 83.6% (**Fig. 3a**). No Citrine2 fluorescence was observed in animals grown on PCC2 but not exposed to UV light or in animals grown without ncAA.

### PC-FLP significantly outperforms PC-Cre in *C. elegans*

We next compared the efficiency of our PC-FLP system to the previously published PC-Cre variants photocaged at the catalytic residues K201 and Y324^9–12^, which correspond to FLP K223 and Y343 respectively.

For a direct comparison, we created transgenic strains equivalent to the PC-FLP strains described above, but expressing PC-Cre. These strains contained the same PylRS/tRNA(M15)_CUA_ variant pairs that we previously used to express PC-FLP, albeit together with Cre variants containing an amber stop codon at K201 or Y324, and a fluorescent reporter with loxP sites flanking the transcriptional stop cassette in place of FRT sites (**Supplementary Fig. 6**).

Animals expressing each of the systems were grown on their respective ncAAs for 24 hours from the L1 larval stage: the tyrosines ONBY, NPY and MNPY, and the lysines PCK and HCK. PC-FLP(C189PCC2) was also included in the assay even though no corresponding cysteine residue exists in Cre. The worms were then illuminated with 365 nm light and Citrine2 expression measured 24 hours later.

We found that for all ncAAs, PC-FLP significantly outperformed PC-Cre. For the photocaged lysines, we measured activation efficiencies in glutamatergic neurons of 83.8% and 77.4% for PC-FLP(K223PCK) and PC-FLP(K223HCK) respectively, compared to 62.3% for PC-Cre(K201PCK) and 56.6% for PC-Cre(K201HCK). Strikingly, all three PC-FLP variants utilising photocaged tyrosine variants, Y343NPY, Y343ONBY and Y343MNPY, showed high efficiencies of 99.1% 99.5% and 98.5% respectively, whereas PC-Cre(Y324NPY), PC-Cre(Y324ONBY), and PC-Cre(Y324MNPY) only yielded efficiencies of 68.9%, 39.2% and 52.4% respectively. PC-FLP(C189PCC2) also out-performed all PC-Cre variants with an efficiency 83.6% (**Fig. 3a**).

The differences became even more striking when we looked at efficiencies at lower ncAA concentrations and reduced the time on ncAA. For these experiments we focused only on the three photocaged tyrosines, which all showed close to 100% activity at 1 mM and 24 hours on ncAA when incorporated in place of Y343 in FLP. When we reduced ncAA concentrations, the best performing PC-FLP variant, PC-FLP(Y343NPY), showed near 100% activity down to 0.01 mM, and 92.5% activity at 0.001 mM. PC-FLP(Y343ONBY) showed 89.3% activity at 0.01 mM and 20.2% at 0.001 mM. In contrast, the best performing PC-Cre(Y324NPY) showed only 2% at 0.01 mM and no detectable activity at 0.001 mM (**Fig. 3b**).

When we reduced the time on ncAA to 8 hours before illumination, thereby reducing the time window for production of PC-FLP and PC-Cre, all three PC-FLP(Y343) variants still showed between 95% to 100% activity at 1 mM, while the activity for all three PC-Cre(Y324) variants was reduced to below 20%. When we reduced the ncAA concentration 100-fold to 0.01 mM, PC-FLP(Y343NPY) animals still showed near 100% activation efficiency and PC-FLP(Y343ONBY) animals showed 82% activation efficiency. In contrast, no PC-Cre animals showed any detectable activity below 0.1 mM (**Fig. 3b**).

### PC-FLP can be used to control expression of target genes with high efficiency and no background

Besides the fluorescent Citrine2 reporter, we also created *C. elegans* lines expressing the different PC-FLP variants in glutamatergic head neurons using the *glr-1p* promoter to control the expression of a Chrimson::F2A::Citrine2 construct that was separated from its promoter by a transcriptional terminator flanked by two FRT sites. When Chrimson is expressed in the glutamatergic head neurons of *C. elegans*, optogenetic activation leads to easily scorable reversals and accelerations^36^. We have previously used PC-Cre to control expression of the optogenetic channels ChR2 and Chrimson in these *C. elegans* neurons, allowing us to control neuronal activity and behaviour^12^.

We grew transgenic animals expressing the PC-FLP systems on 1 mM of the corresponding photocaged ncAA for 48 hours from the L1 larval stage. We illuminated the animals with 365 nm light and then transferred them to NGM agar plates supplemented with all-trans retinal (ATR) for 24 hours to give time for expression of the functional Chrimson channel. We then performed optogenetic behavioural assays by illuminating the animals for 1 second using a 617 nm LED to activate Chrimson. Activity was scored by determining the fraction of animals responding to the stimulus.

In animals expressing PC-FLP, photocaged with either the photocaged lysine PCK or the photocaged tyrosines NPY, ONBY, MNPY, we found that 100% of animals responded to the stimulus. In the strains expressing FLP photocaged with PCC2, 90% of animals responded to the stimulus (**Fig. 3c**). Importantly, we saw no response in animals grown on any of the ncAAs, but not illuminated, further confirming that all photocaged ncAAs fully block the function of FLP and allow tight control of target gene expression.

Next, we explored whether PC-FLP can be used to control gene expression at a genomic locus. For this, we used a strain that contains a genomically integrated reporter consisting of nuclear localised mCherry::his-58 flanked by FRT sites followed by GFP::his-58, driven by the *hsp-16.41* heat shock promoter^4^. This promoter is activated after heat shock in around 50% of *C. elegans* neurons^37^. Without FLP activity, activation of the promoter by heat shock results in the expression of mCherry alone. In the presence of active FLP, the mCherry gene and 3’UTR are excised, resulting in expression of GFP alone. Since each cell contains two copies of the reporter locus, low FLP activity could result in expression of both mCherry and GFP. Based on this strain, we generate transgenic strains expressing the NPYRS/tRNA(M15)_CUA_ pair and FLP containing an amber stop codon at position Y343 in an operon with nuclear localised NLS::mTagBFP2 **(Supplementary Fig. 7a)**. Expression of PylRS and FLP variant was driven by the promoter for the glutamate receptor *glr-1* that is active in a subset of *C. elegans* head neurons^12^.

These Transgenic animals were then grown on NGM plates supplemented with 1 mM NPY to express PC-FLP(Y343NPY). To assay PC-FLP(Y343NPY) activity, the animals were then either illuminated with 365 nm light or kept in the dark, and reporter gene expression was induced by heat shock 12h later.

We then assayed cells expressing PC-FLP, as indicated by the presence of mTagBFP2 fluorescence, for expression of the reporter. In illuminated animals, we found only GFP fluorescence in these cells, but no mCherry fluorescence, indicating complete recombination of the genomic reporter locus **(Fig. 3d, Supplementary Fig. 7b)**. In contrast, when we looked at animals that were kept in the dark, we instead found only cells expressing mCherry, and no cells expressing GFP, confirming that PC-FLP(Y343NPY) is indeed fully caged and does not display any residual background activity.

### PC-FLP is functional in tissues outside the nervous system

We next wanted to determine whether PC-FLP could be generally applied in tissues other than the nervous system. For this, we generated strains to express the best performing PC-FLP variant, PC-FLP(Y343NPY), in either the pharynx, or body wall muscle cells. The synthetase NPYRS and FLP were driven by the *myo-2p* promoter for expression in the pharynx, or the *myo-3p* promoter for expression in body wall muscle cells. In both cases we used the ubiquitous *eef-1A.1p* (*eft-3p)* promoter to drive expression of the Citrine2 reporter gene.

Transgenic animals were grown from the L1 larval stage for 48 hours on NGM agar plates containing 1 mM NPY to express PC-FLP(Y343NPY), followed by illumination at 365 nm. 24 hours after illumination worms were assayed for Citrine2 fluorescence. In both cases for animals expressing PC-FLP(Y343NPY) in either the pharynx or the body wall muscle, we observed clearly visible Citrine2 fluorescence. In contrast, we observed no fluorescence in animals grown on 1 mM NPY but not illuminated (**Fig. 4a, b**).

**Fig. 4.**
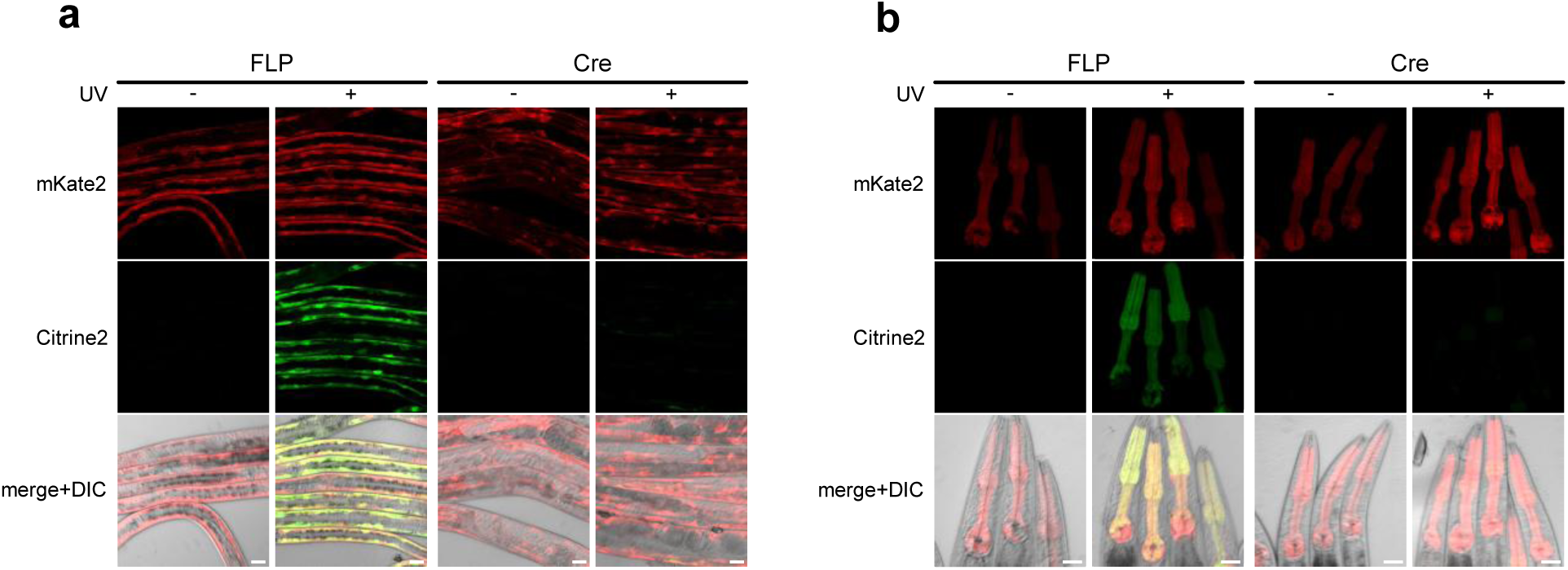
PC-FLP and PC-Cre in the body wall muscle and the pharynx. **a** Fluorescence microscopy of worms expressing the PC-FLP and PC-Cre systems using the photocaged tyrosine NPY in the body wall muscle driven by the promoter for *myo-3*. Worms were grown in the presence of NPY at a concentration of 1 mM for 48 hours and then illuminated with 365 nm ("+" condition) or kept in the dark ("-" condition). Animals were imaged 24 hours after illumination. Scale bars represent: 20 µm. **b** Fluorescence microscopy of worms expressing the PC-FLP and PC-Cre systems using the photocaged tyrosine NPY in the pharynx driven by the promoter for *myo-2*. Worms were grown in the presence of NPY at a concentration of 1 mM for 48 hours and then illuminated with broad-band UV light ("+" condition) or kept in the dark ("-" condition). The worms were then imaged 24 hours after illumination. Scale bars represent: 20 µm.

To further compare FLP to Cre, we also generated strains expressing the corresponding PC-Cre(Y324NPY) in the pharynx and in body wall muscle. Here, in contrast to PC-FLP(Y343NPY), we found no strains that displayed more than very weak Citrine2 expression in few cells for both tissues, showing that PC-FLP enables optical control of gene expression in tissues inaccessible to PC-Cre (**Fig. 4a, b, Supplementary Fig. 8**).

### PC-FLP can be efficiently activated with wavelengths longer than 36 nm

One of the main advantages of controlling the activity of SSRs using photocaged ncAAs is the speed of the uncaging reaction allowing for the permanent activation of the SSR using a very brief pulse of light in the low second or even sub-second range. This light can be targeted very precisely to localise the activation to single cells in living animals^12^. We have previously carried out single-cell activation of PC-Cre using a microscope-mounted laser to deliver 365 nm light. While the photocaging groups used here are most sensitive to wavelengths below 400 nm, we nevertheless wanted to test activation efficiencies at longer wavelengths, since it would be highly advantageous to be able to efficiently activate PC-FLP at a wavelength of 405 nm so that single cell activation could be carried out using a standard 405 nm laser commonly used on confocal microscopes.

When we recorded the absorption spectra of the photocaged ncAAs used here, we found that, encouragingly, all ncAAs absorbed at 405 nm, albeit with ONBY showing lower absorption than the other compounds (**Supplementary Fig. 9a**).

We first assayed the activation of PC-FLP at different wavelength ranges using LED illumination. For this, animals expressing the PC-FLP systems were grown from the L1 larval stage for 24 hours on NGM agar plates containing their respective ncAA at a concentration of 1 mM. The worms were then illuminated for 5 minutes using LEDs to deliver light with peak wavelengths of 365 nm, 405 nm, 435 nm, 460 nm, or 470 nm (**Supplementary Fig. 9b**). In all cases, we found that PC-FLP variants activated efficiently at 365 nm, and that there was no significant difference between the activation efficiency with the 365 nm LED and 405 nm LED. With the 435 nm LED, we observed none or only marginal activation of PC-FLP in animals expressing PC-FLP(K223PCK), PC-FLP(K223HCK), and PC-FLP(Y343ONBY), while animals expressing PC-FLP(Y343NPY) showed an activation efficiency of 6%. In contrast, animals expressing PC-FLP(Y343MNPY) or PC-FLP(C189PCC2) both showed efficient activation of 55% and 68.8% respectively (**Fig. a**).

Importantly, we further tested the stability of the PCC2 and MNPY photocages at wavelengths above 435 nm and found that there was only negligible activation of approximately 1% for PC-FLP(C189PCC2) with the 460 nm LED, and no activation with the 470 nm LED. PC-FLP(Y343MNPY) was not activated by either 460 nm or 470 nm LED illumination. Furthermore, we observed no activation after illuminating using a standard 480/20 nm GFP filter for 5 minutes or exposing animals expressing PC-FLP(C189PCC2) to standard laboratory ambient lighting for 24 hours, demonstrating that PC-FLP(C189PCC2) and PC-FLP(Y343MNPY) are still compatible with blue-light optogenetics and imaging (**Supplementary Fig. 9c**).

### Activation of PC-FLP using a 4 nm microscope mounted laser

We then proceeded to test activation of the different PC-FLP variants using a standard confocal 405 nm laser. For this, animals were again grown from the L1 larval stage for 48 hours on NGM agar plates containing their respective ncAA at a concentration of 1 mM. They were then anaesthetized, mounted on agar slides, and we scanned their head region using the 405 nm laser at various power settings. The animals were then recovered onto NGM agar plates and imaged to quantify activation efficiency 24 hours after illumination (**Fig. b, Supplementary Fig. 9d**). We found that at 50% laser power, PC-FLP(Y343NPY), PC-FLP(Y343MNPY), and surprisingly even PC-FLP(Y343ONBY) were activated with near 100% efficiency. At 10% laser power, PC-FLP(Y343NPY) was still activated with near 100% efficiency, and PC-FLP(Y343MNPY) with 84.3% efficiency, but we saw no activation of PC-FLP(Y343ONBY). At 1% laser power we only saw activation of PC-FLP(Y343MNPY) at 8.3% efficiency. For the PC-FLP variants containing the photocaged lysines, PC-FLP(K223PCK) and PC-FLP(K223HCK), at 50% laser power we observed activation efficiencies of 33.4% and 21.4% respectively. At 10% laser power we saw only minimal activation of PC-FLP(K223PCK) (1.9%), and no activity for PC-FLP(K223HCK). Interestingly, PC-FLP(C189PCC2) never displayed activity above 64.5%, but showed similar activation efficiencies at all three power levels of 50%, 10% and 1% respectively.

Having confirmed that several PC-FLP variants can be efficiently activated using a 405 nm laser at a range of power levels, we next tested whether the confocal microscope would serve to precisely target PC-FLP activation to single cells. We carried out this experiment with the PC-FLP(Y343NPY) animals at 10% laser power and targeted the bilaterally symmetric PVC neurons in the tail, since the *glr-1p* promoter used to drive FLP and NPYRS is active in these neurons.

Worms were grown from the L1 larval stage for 48 hours on NGM agar plates containing 1 mM NPY. They were then anaesthetized and mounted on agar slides. We then targeted the laser to one of the PVCs using the bleaching function to illuminate in a single Z-stack, with a total illumination time per cell of below 10 ms. 24 hours after illumination we observed Citrine2 only in the cell that was targeted by the laser (Fig. 5c).

**Fig. 5.**
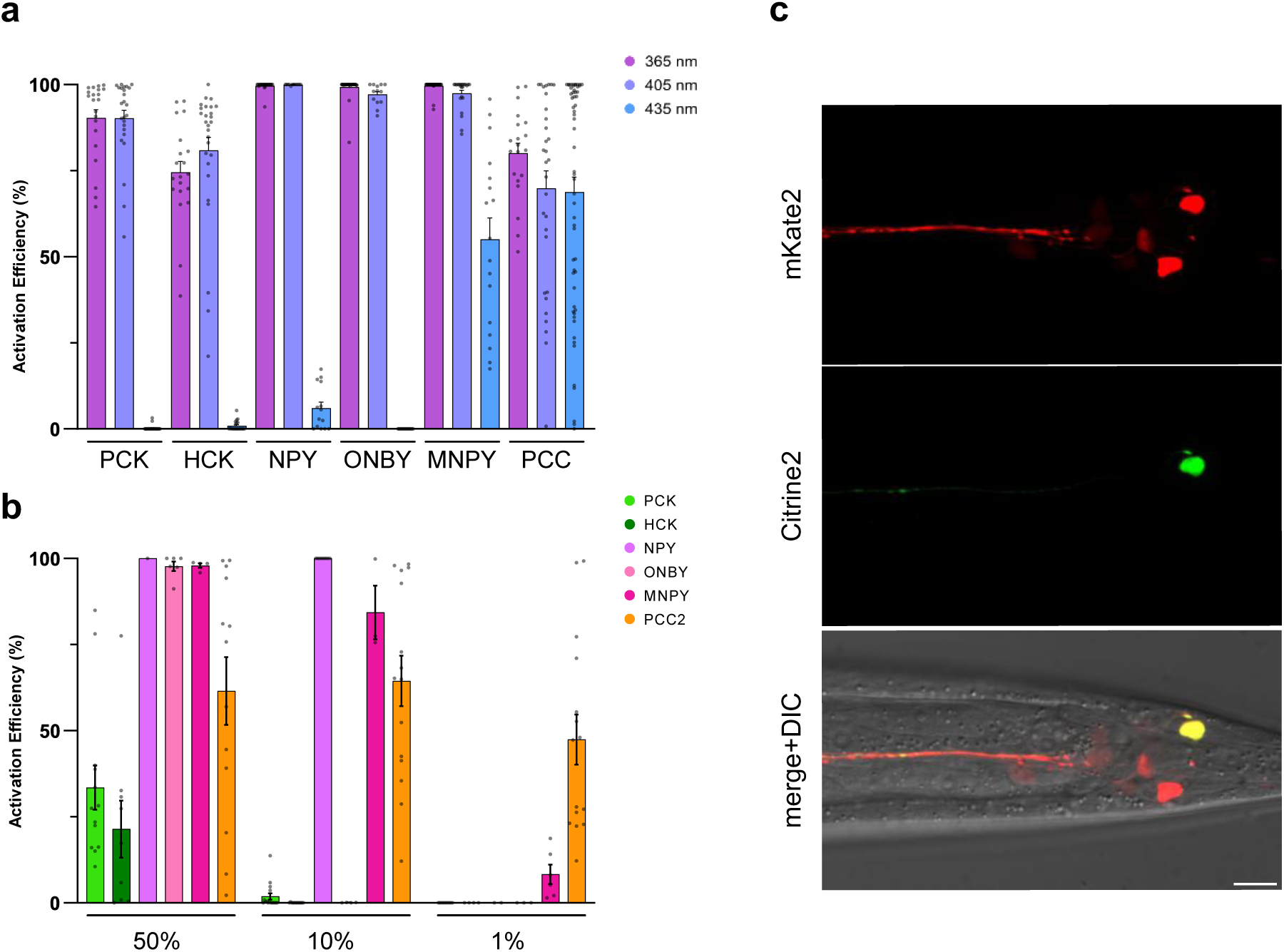
Activation of PC-FLP variants at longer wavelengths and targeted activation using a 4 nm laser. **a** Activation efficiency in animals expressing PC-FLP using the different photocaged variants of tyrosine, lysine, and cysteine following illumination using LEDs of various wavelengths. Worms were grown in the presence of the respective ncAA at a concentration of 1 mM for 24 hours and then illuminated for 5 minutes. The worms were then imaged for quantification 24 hours after illumination. Dots represent individual animals assayed. Error bars represent the SEM. **b** Activation efficiency in animals expressing PC-FLP using the different photocaged variants of tyrosine, lysine, and cysteine following illumination with the 405 nm laser on the Zeiss LSM800 confocal microscope. Worms were grown in the presence of the respective ncAA at a concentration of 1 mM for 48 hours and then illuminated for 5 minutes. The worms were then imaged for quantification 24 hours after illumination. Percentage categories represent the pseudo-percentage laser power settings. Dots represent individual animals assayed. Error bars represent the SEM. **c** Fluorescence microscopy images showing expression of Citrine2 in a single PVC neuron 24 hours after targeted illumination with the 405 nm laser on the Zeiss LSM800. Scale bar represents: 10 µm.

## DISCUSSION

We develop a photoactivatable FLP recombinase by incorporating photocaged non-canonical amino acids into FLP using genetic code expansion technology. We demonstrate that the activity of FLP can be photocaged by replacing any one of the active site residues Tyr343, Lys223, and Cys189 with their respective photocaged counterpart. We find that in all cases, PC-FLP variants function more efficiently than PC-Cre and that in general, PC-FLP outperforms PC-Cre under all conditions and in all tissues we have tested.

We further establish that PC-FLP can be efficiently activated using light with wavelengths longer than 365 nm, thus allowing the use of a standard 405 nm microscope-mounted laser to activate the expression of target genes with single-cell precision. This makes PC-FLP easily accessible for most laboratories as a tool to control gene expression with single cell precision, without the need for specialised equipment. The short illumination time of less than 10 ms per cell that is required to activate PC-FLP provides what is essentially an easy and effective "point-and-click" method to control gene expression in an animal.

While PC-FLP can be activated by light between 365 to 435 nm, importantly, all variants we tested remain entirely inactive at wavelengths above 450nm, thus making them fully compatible with most commonly used imaging and optogenetic approaches.

Based on the systematic characterisation of photocaged FLP and Cre variants presented here, we can make recommendations for users wishing to apply them for their research. While the most efficient variant is PC-FLP(Y343NPY), we suggest that PC-FLP(Y343ONBY) may be a better starting point for most users, since it uses the commercially available and inexpensive tyrosine variant ONBY, whereas NPY is currently only available via custom synthesis. FLP(Y343ONBY), albeit only the second-best performing variant at 365 nm, is still highly efficient and suitable for activation using a 405nm microscope mounted laser.

For ncAAs that are not currently commercially available, detailed synthesis protocols can be found in the relevant literature^22–24,26^ and the required precursors are commercially available. Any academic organic chemistry laboratory or company offering custom chemical synthesis should therefore be able to easily synthesise the compounds. For commercial availability of ncAAs used in this study see also Supplementary Note 2.

Looking ahead, the availability of two SSRs, photocageable using various photocaged ncAAs (Lys, Tyr, Cys) raises the exciting future possibility of combining PC-FLP and PC-Cre, photocaged by different ncAAs, to create even more powerful tools for optically controlling gene expression. Our finding that some of the photocaged ncAAs tested (MNPY, PCC2) can be efficiently uncaged at longer wavelengths, while others show no uncaging at these wavelengths (PCK, HCK) suggests the prospect of differentially activating PC-FLP and PC-Cre using different wavelengths of light.

The use of two photocaged ncAAs would however require mutually orthogonal PylRS/tRNA pairs for the incorporation of one photocaged ncAA into FLP and a different photocaged ncAA into Cre in the same cell. While several such mutually orthogonal PylRS/tRNA(Pyl) pairs have been established in *E. coli* and mammalian cell culture systems^38–41^, none have thus far been reported for multicellular organisms.

Additionally, apart from the PylRSs from *Methanosarcina* species, which are not mutually orthogonal, no PylRSs are available for incorporation of photocaged ncAAs. It is however foreseeable that these limitations will be overcome in the near future, opening the path towards the use of two individually controllable photocaged SSRs in the same cell.

The fact that FLP activity can be precisely controlled through the incorporation of photocaged ncAAs suggests the possibility that this approach will be transferrable to other SSRs beside FLP and Cre. Several other tyrosine SSRs related to FLP and Cre, such as Dre, Vika, B3, KD, have been established in a variety of different models. Given the high sequence homology to some members and conserved mechanism of action, it is very likely that these too could be photocaged by replacing critical amino acid residues with photocaged variants as outlined here.

The ncAAs we have employed here are light sensitive in the 365nm to 435 nm range, and stable at longer wavelengths. While this has the advantage of making them compatible for use together with most other optogenetic and imaging approaches, such shorter uncaging wavelengths are not well suited for deep tissue penetration. An advantage of the genetic code expansion approach is however that it should be easily possible to directly install other ncAA with different uncaging properties into PC-FLP and PC-Cre in the future. Already available examples of such ncAA are lysine derivatives that are green light sensitive at 520 nm^42^, or amenable to two-photon uncaging at 730 nm^23^. Both these photo-caged lysines utilise the *M. mazei* PylRS/tRNA(Pyl) system we employ here.

While this work has been carried out in *C. elegans*, the components it uses are not specific to worms and may therefore be transferable to other systems in which FLP and genetic code expansion have been established, including bacteria, cultured mammalian cells and organoids, as well as other animal models such zebrafish^43^, *Drosophila*^44^, and mouse^45^.

## DATA AVAILABILIT

All data are included in the manuscript and the supplementary file. Sequences that have been used for transgenic constructs are listed in the supplementary file.

## ACKNOWLEDGEMENTS

We thank Maria Doitsidou, Deep Prakash and Rebekah White for helpful suggestions on the manuscript. We thank Peter Askjaer for strain BN294. We are grateful to Maria Doitsidou for assistance with imaging, and to the staff at the IMPACT imaging facility at the University of Edinburgh for assistance in the initial setup of laser uncaging experiments.

## AUTHOR CONTRIBUTIONS

J.J.V.R and S.G. designed experiments, J.J.V.R performed experiments and analysed data, K.B. performed initial experiments, Y.M. performed heat shock experiments and assisted with some imaging, J.J.V.R and S.G wrote the manuscript.

## FUNDING

The work was supported by the European Research Council [ERC-StG-679990], the BBSRC [BB/Y006380/1 & BB/W014610/1], The Royal Society, The Muir Maxwell Epilepsy Centre.

## Notes

### Competing Interest Statement

The authors have declared no competing interest.

### Summary of Updates

We have carried out additional experiments to demonstrate control of gene expression using PC-FLP at an genomic locus. We have also added further details on how recombinase activity was quantified, and validated our automated analysis with an experiment in which it was compared to directly counting activated cells.

## REFERENCES

1. Fischer, J. A., Giniger, E., Maniatis, T. & Ptashne, M. GAL4 activates transcription in Drosophila. Nature 332, 853–856 (1988).

2. Meinke, G., Bohm, A., Hauber, J., Pisabarro, M. T. & Buchholz, F. Cre Recombinase and Other Tyrosine Recombinases. Chem. Rev. 116, 12785–12820 (2016).

3. Feil, R. et al. Ligand-activated site-specific recombination in mice. Proc. Natl. Acad. Sci. 93, 10887–10890 (1996).

4. Muñoz-Jiménez, C. et al. An Efficient FLP-Based Toolkit for Spatiotemporal Control of Gene Expression in *Caenorhabditis elegans*. Genetics 2 **6**, 1763–1778 (2017).

5. Schmitt, C., Schultheis, C., Husson, S. J., Liewald, J. F. & Gottschalk, A. Specific Expression of Channelrhodopsin-2 in Single Neurons of Caenorhabditis elegans. PLoS ONE 7, e43164 (2012).

6. Jung, H. et al. Noninvasive optical activation of Flp recombinase for genetic manipulation in deep mouse brain regions. Nat. Commun. 1 , 314 (2019).

7. Schindler, S. E. et al. Photo-activatable Cre recombinase regulates gene expression in vivo. Sci. Rep. 5, 13627 (2015).

8. Charette, M., Rosenblum, C., Shade, O. & Deiters, A. Optogenetics with Atomic Precision─A Comprehensive Review of Optical Control of Protein Function through Genetic Code Expansion. Chem. Rev. 12 , 1663–1717 (2025).

9. Edwards, W. F., Young, D. D. & Deiters, A. Light-Activated Cre Recombinase as a Tool for the Spatial and Temporal Control of Gene Function in Mammalian Cells. ACS Chem. Biol. 4, 441–445 (2009).

10. Luo, J. et al. Genetically encoded optical activation of DNA recombination in human cells. Chem. Commun. 2, 8529–8532 (2016).

11. Brown, W., Liu, J., Tsang, M. & Deiters, A. Cell-Lineage Tracing in Zebrafish Embryos with an Expanded Genetic Code. ChemBioChem 19, 1244–1249 (2018).

12. Davis, L. et al. Precise optical control of gene expression in C. elegans using improved genetic code expansion and cre recombinase. eLife 1 , 1–22 (2021).

13. Broach, J. R. & Hicks, J. B. Replication and Recombination Functions Associated with the Yeast Plasmid, 2μ Circle. Cell 21, 501–508 (1980).

14. Buchholz, F. Different thermostabilities of FLP and Cre recombinases: implications for applied site-specific recombination. Nucleic Acids Res. 24, 4256–4262 (1996).

15. Buchholz, F., Angrand, P.-O. & Stewart, A. F. Improved properties of FLP recombinase evolved by cycling mutagenesis. Nat. Biotechnol. 16, 657–662 (1998).

16. Heidmann, D. & Lehner, C. F. Reduction of Cre recombinase toxicity in proliferating Drosophila cells by estrogen-dependent activity regulation. Dev. Genes Evol. 211, 458–465 (2001).

17. Nern, A., Pfeiffer, B. D., Svoboda, K. & Rubin, G. M. Multiple new site-specific recombinases for use in manipulating animal genomes. Proc. Natl. Acad. Sci. 1 **8**, 14198–14203 (2011).

18. Brenner, S. The genetics of Caenorhabditis elegans. Genetics 77, 71–94 (1974).

19. Stiernagle, T. Maintenance of C. elegans (February 11, 2006). in Wormbook (ed. Community, T. C. elegans R.) (WormBook, 2006).

20. Radman, I., Greiss, S. & Chin, J. W. Efficient and rapid C. elegans transgenesis by bombardment and hygromycin B selection. PLoS One 8, e76019 (2013).

21. Greiss, S. & Chin, J. W. Expanding the Genetic Code of an Animal. J. Am. Chem. Soc. 133, 14196–14199 (2011).

22. Gautier, A. et al. Genetically encoded photocontrol of protein localization in mammalian cells. J. Am. Chem. Soc. 132, 4086–4088 (2010).

23. Luo, J. et al. Genetically encoded optochemical probes for simultaneous fluorescence reporting and light activation of protein function with two-photon excitation. J. Am. Chem. Soc. 136, 15551–15558 (2014).

24. Luo, J., Torres-Kolbus, J., Liu, J. & Deiters, A. Genetic Encoding of Photocaged Tyrosines with Improved Light-Activation Properties for the Optical Control of Protease Function. ChemBioChem 18, 1442–1447 (2017).

25. Deiters, A., Groff, D., Ryu, Y., Xie, J. & Schultz, P. G. A genetically encoded photocaged tyrosine. Angew. Chem. 118, 2794–2797 (2006).

26. Nguyen, D. P. et al. Genetic Encoding of Photocaged Cysteine Allows Photoactivation of TEV Protease in Live Mammalian Cells. J. Am. Chem. Soc. 136, 2240–2243 (2014).

27. Crnković, A., Suzuki, T., Söll, D. & Reynolds, N. M. Pyrrolysyl-tRNA Synthetase, an Aminoacyl-tRNA Synthetase for Genetic Code Expansion. Croat. Chem. Acta 89, (2016).

28. Nunes-Duby, S. E., Kwon, H. J., Tirumalai, R. S., Ellenberger, T. & Landy, A. Similarities and differences among 105 members of the Int family of site-specific recombinases. Nucleic Acids Res. 26, 391–406 (1998).

29. Chen, J.-W., Evans, B. R., Yang, S.-H., Araki, H. & Jayaram, M. Functional Analysis of Box I Mutations in Yeast Site-Specific Recombinases Flp and R: Pairwise Complementation with Recombinase Variants Lacking the Active-Site Tyrosine. MOL CELL BIOL 12, (1992).

30. Xi, Z. et al. Using a quadruplet codon to expand the genetic code of an animal. Nucleic Acids Res. , 4801–4812 (2022).

31. Nikić, I., et al. Debugging Eukaryotic Genetic Code Expansion for Site-Specific Click-PAINT Super-Resolution Microscopy. Angew. Chem. Int. Ed., 16172–16176 (2016).

32. Serfling, R. et al. Designer tRNAs for efficient incorporation of non-canonical amino acids by the pyrrolysine system in mammalian cells. Nucleic Acids Res. 46, 1–10 (2018).

33. O’Shea, J. M. et al. Generation of Photocaged Nanobodies for Intracellular Applications in an Animal Using Genetic Code Expansion and Computationally Guided Protein Engineering**. ChemBioChem 23, e202200321 (2022).

34. Tseng, H.-W. et al. Expanding the Scope of Orthogonal Translation with Pyrrolysyl-tRNA Synthetases Dedicated to Aromatic Amino Acids. Molecules 2 , 4418 (2020).

35. Chen, Y. Crystal Structure of a Flp Recombinase–Holliday Junction Complex Assembly of an Active Oligomer by Helix Swapping. Mol. Cell 6, 885–897 (2000).

36. Stirman, J. N. et al. Real-time multimodal optical control of neurons and muscles in freely behaving Caenorhabditis elegans. Nat. Methods 8, 153–158 (2011).

37. Stringham, E. G., Dixon, D. K., Jones, D. & Candido, E. Temporal and spatial expression patterns of the small heat shock (hsp16) genes in transgenic Caenorhabditis elegans. Mol. Biol. Cell 3, 221–233 (1992).

38. Willis, J. C. & Chin, J. W. Mutually orthogonal pyrrolysyl-tRNA synthetase/tRNA pairs. Nat. Chem. 1 , 831–837 (2018).

39. Beránek, V., Willis, J. C. & Chin, J. W. An evolved Methanomethylophilus alvus pyrrolysyl-tRNA synthetase/tRNA pair is highly active and orthogonal in mammalian cells. Biochemistry 8, 387–390 (2018).

40. Meineke, B., Heimgärtner, J., Lafranchi, L. & Elsässer, S. J. Methanomethylophilus alvus Mx1201 provides basis for mutual orthogonal pyrrolysyl tRNA/aminoacyl-tRNA synthetase pairs in mammalian cells. ACS Chem. Biol. 13, 3087–3096 (2018).

41. Meineke, B., Heimgärtner, J., Eirich, J., Landreh, M. & Elsässer, S. J. Site-specific incorporation of two ncAAs for two-color bioorthogonal labeling and crosslinking of proteins on live mammalian cells. Cell Rep. 31, 107811 (2020).

42. Li, M. et al. Genetically Encoded Green-light-responsive Photocaged Lysine for Sequential Control of Protein Function. Chem. Sci. (2025).

43. Liu, J., Hemphill, J., Samanta, S., Tsang, M. & Deiters, A. Genetic code expansion in zebrafish embryos and its application to optical control of cell signaling. J. Am. Chem. Soc. 139, 9100–9103 (2017).

44. Bianco, A., Townsley, F. M., Greiss, S., Lang, K. & Chin, J. W. Expanding the genetic code of Drosophila melanogaster. Nat. Chem. Biol. 8, 748–750 (2012).

45. Chen, Y. et al. Heritable expansion of the genetic code in mouse and zebrafish. Cell Res. 27, 294–297 (2017).

