## Supplementary for "Photocaged FLP Recombinase for Precise Spatio-temporal Control of Gene Expression"

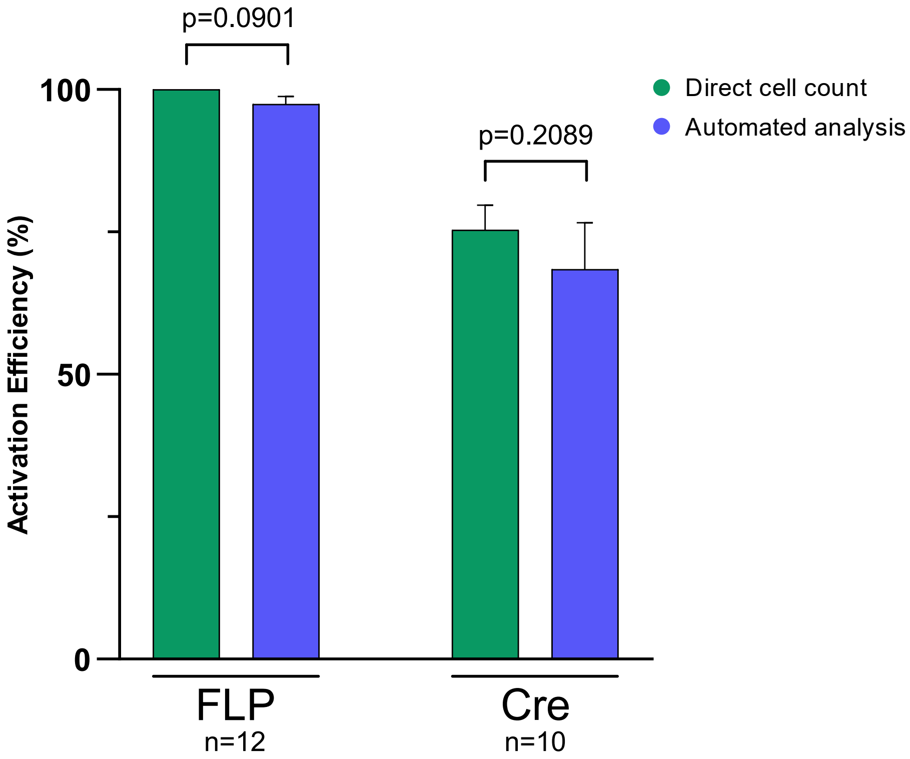


**Supplementary Fig. 1 Comparison of automated analysis and direct cell count for quantification of recombination.** Activation efficiency of PC-FLP(Y343NPY) and PC-Cre(Y324NPY) following illumination with 365 nm light determined by direct cell count or by automated analysis. Then same images of animals were assayed once by direct counting and then by the automated analysis pipeline. For the direct count the value graphed represents the percentage of cells expressing either PC-FLP or PC-Cre as indicated by mKate2, that also express the reporter Citrine2. For the automated count the value graphed represents the percentage of mKate2 expressing area that also expresses Citrine2. Error bars represent the SEM. p values correspond to a paired t-test, and n values represent the number of animals that were analysed by both approaches.

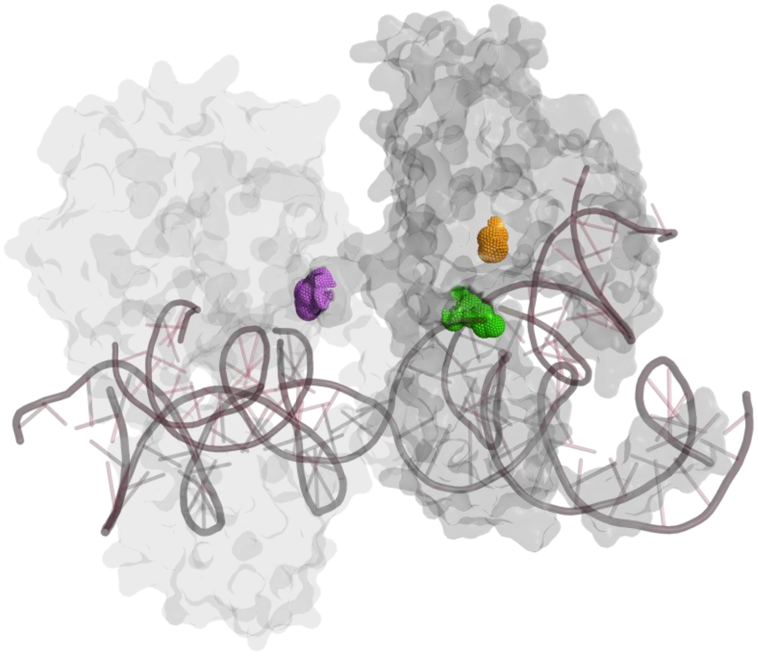

**Supplementary Fig. 2** Overview of the candidate positions to photocage the activity of FLP. On the FLP molecule on the right side, K223 and C189, which act in *cis*, are highlighted in green and orange respectively. Y343, which forms the active site with another FLP molecule (lighter grey on the left side) in trans, is highlighted in purple (PDB: 1FLO[1]).

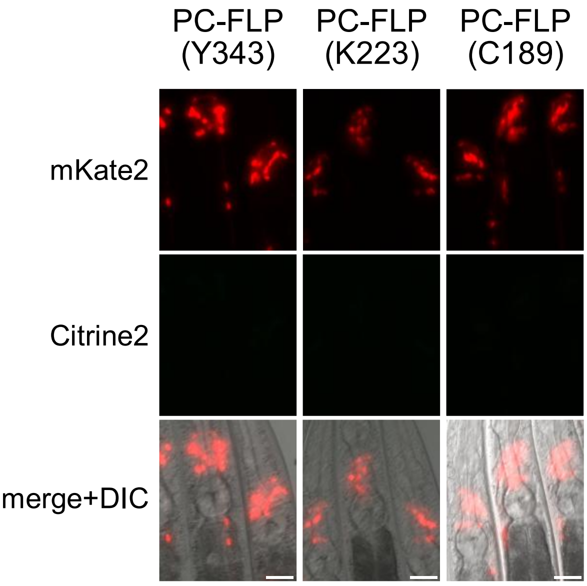


**Supplementary Fig. 3 PC-FLP in the absence of ncAA.** Fluorescence microscopy of worms expressing the PC-FLP(Y343), PC-FLP(K201), or PC-FLP(C189) system grown in the absence of ncAA. There is no expression of the Citrine2 reporter. Scale bars represent: 20 μm.

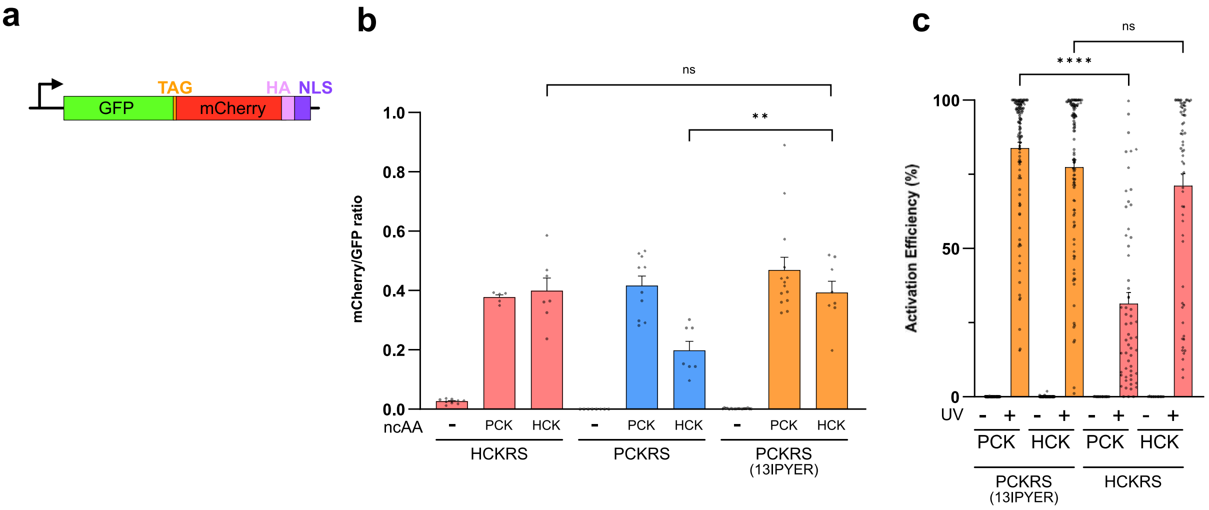


**Supplementary Fig. 3 Comparison of HCKRS and PCKRS. a** Schematic of GFP::mCherry construct used to quantify incorporation by the various PylRS variants. **b** mCherry/GFP ratio of animals expressing HCKRS, PCKRS, or PCKRS(13IPYER) grown for 24 hours on plates with no added ncAA ("-" condition), 1 mM of PCK, or 1 mM of HCK. Dots represent groups of animals assayed. Error bars represent the SEM. Significance values are determined by Welch's t-test. **c** Activation efficiency of animals expressing the HCKRS or PCKRS(13IPYER) to incorporate PCK or HCK into FLP recombinase. Worms were grown in the presence of the 1 mM PCK or 1 mM HCK for 24 hours and then illuminated for 5 minutes with 365 nm light ("+" condition) or kept in the dark ("-" condition). The worms were then imaged for quantification 24 hours after illumination. Dots represent individual animals assayed. Error bars represent the SEM. Significance values are determined by Welch's t-test.

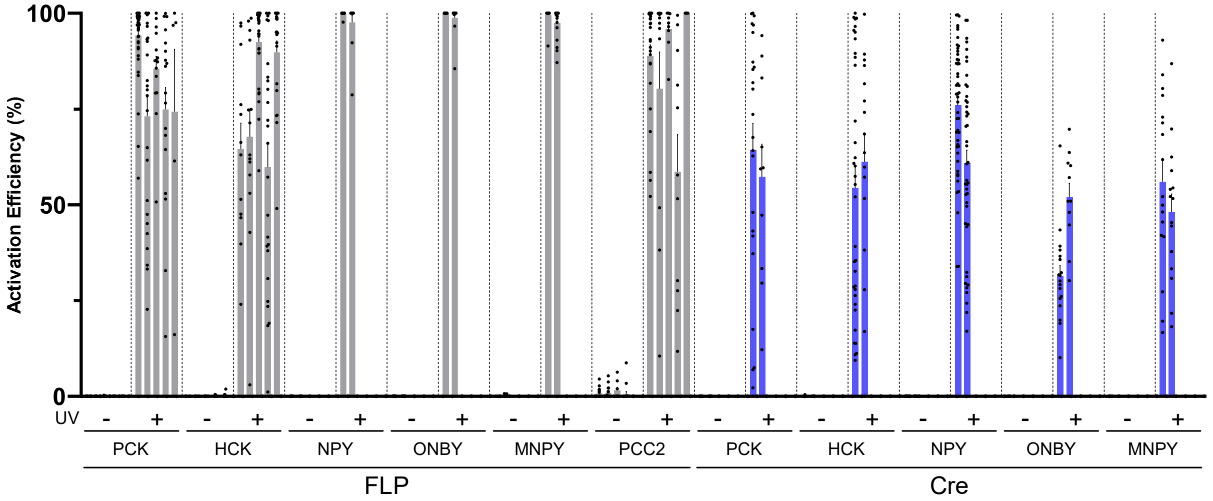


**Supplementary Fig. 5 Activation efficiency measurements for individual independent transgenic strains**. Individual bars in each condition represent separate transgenic strains assayed for Figure 3 a. Error bars represent the SEM.

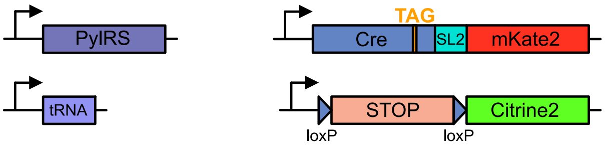


**Supplementary Fig. 6 Expression of PC-Cre.** Constructs transformed into *C. elegans* to assay PC-Cre activity. Equivalent to the constructs used to assay PC-FLP activity.

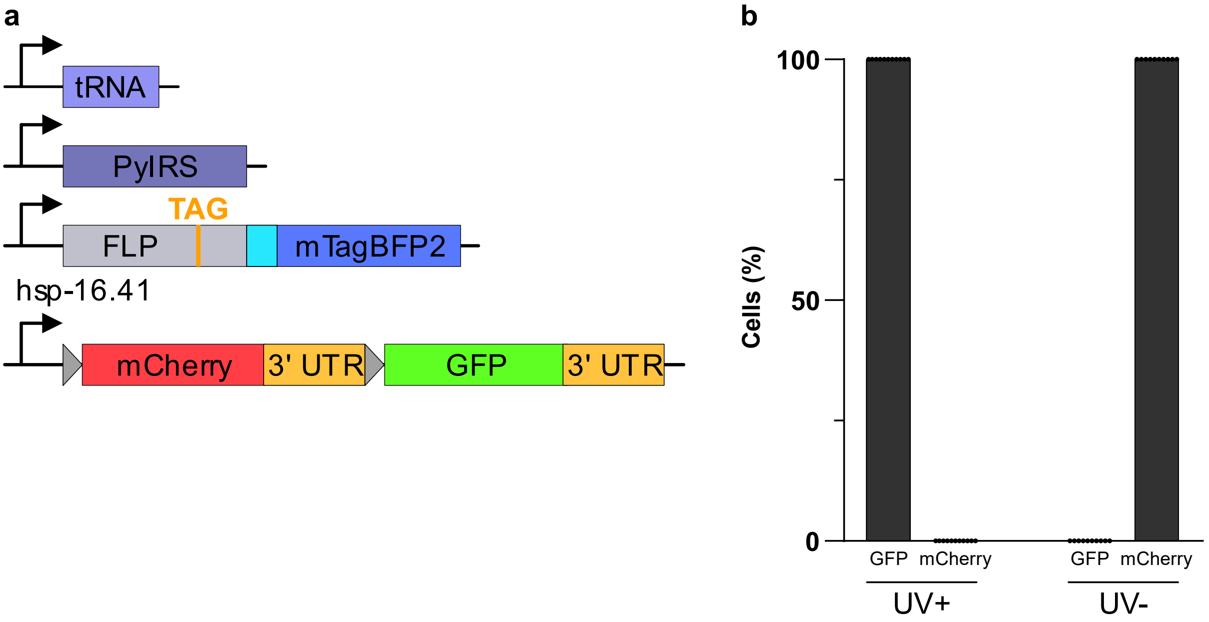


**Supplementary Fig. 7 Optical control of recombination at a genomic locus. a** Constructs transformed into *C. elegans* to express PC-FLP(Y343NPY), and genomically integrated reporter consisting of mCherry flanked by FRT sites upstream of GFP. **b** Percentage of cells expressing either GFP or mCherry fluorescence after heat shock. We did not consider cells where the heat shock promoter *hsp-16.41* is not active, as evident by the lack of either mCherry or GFP expression, and cells that did not express PC-FLP (mTagBFP2). Data correspond to two experimental repeats, consisting of a total of 11 animals (UV+) and 10 animals (UV-).

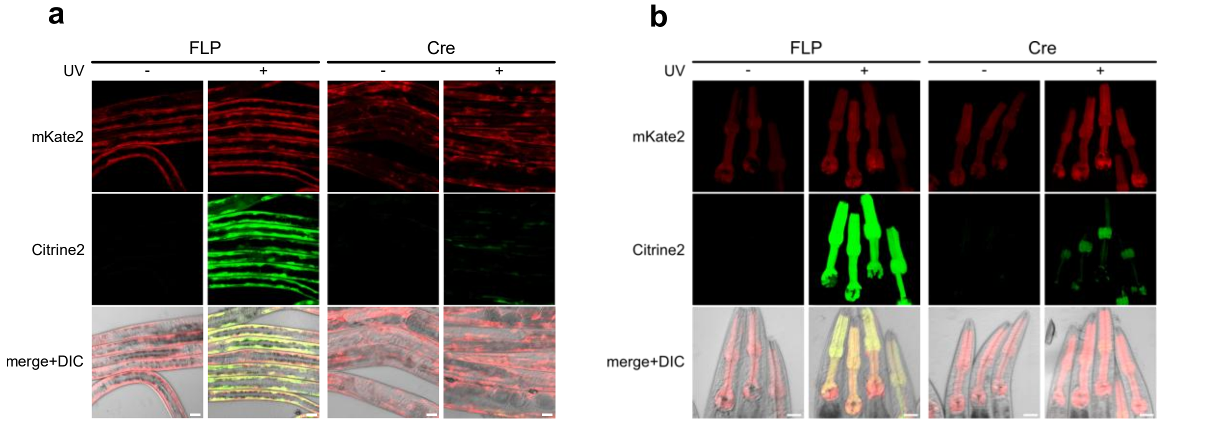


**Supplementary Fig. 8 PC-FLP and PC-Cre in the body wall muscle and the pharynx.** Fluorescence microscopy of worms expressing PC-FLP(Y343NPY) or PC-Cre(Y324NPY) in the body-wall or pharynx. Same set of images as in **Figure 4** but with brightness in the Citrine2 channel increased by **a** 100%, **b** 200% for clear visualization of inferior PC-Cre activity.

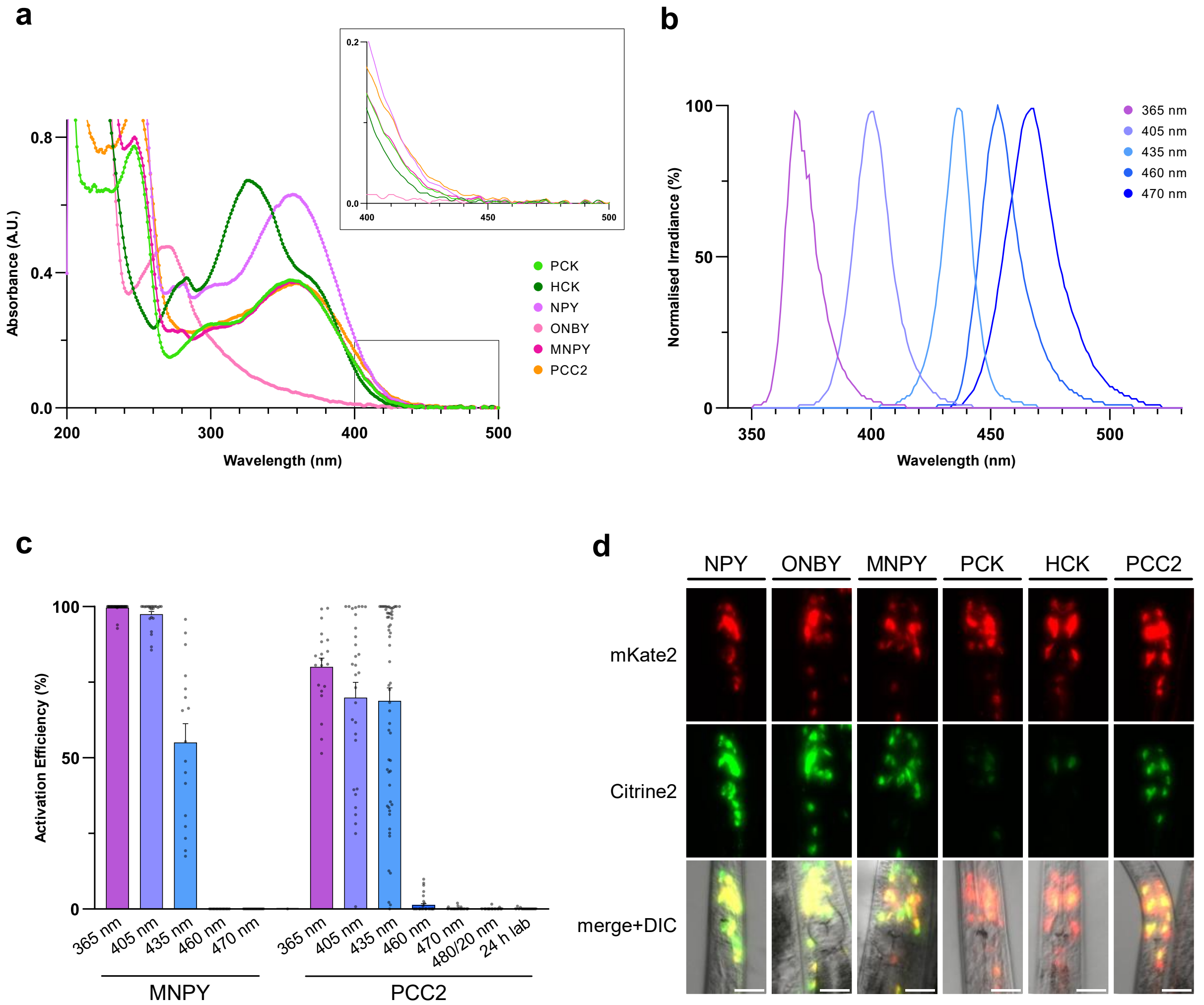


**Supplementary Fig. 9 Activation of PC-FLP variants at longer wavelengths. a** UV-vis absorbance of the different photocaged variants of tyrosine, lysine, and cysteine. **b** Relative irradiance of the "365 nm", "405 nm", "435 nm", "460 nm", "470 nm" LEDs from the pE-4000 light source (CoolLED). Spectra sourced from FPbase[2]. **c** Activation efficiency of animals expressing PC-FLP(Y343MNPY) and PC-

FLP(C189PCC2) 24 hours after illumination by “365 nm”, “405 nm”, “435 nm”, “460 nm”, “470 nm” LEDs from the pE-4000 light source (CoolLED) for 5 minutes. Additionally, activation efficiency of animals expressing PC-FLP(C189PCC2) that were exposed to the output of the HXP120 light source (Leistungselektronik Jena) through a 480/20 nm filter (Oxford Instruments Andor, MM-7253-GFP), and of worms that were left exposed to laboratory fluorescent lights for 24 hours. **d** Fluorescence microscopy of worms expressing the PC-FLP system using the different photocaged variants of tyrosine, lysine, and cysteine and activated using the 405 nm laser on a Zeiss LSM800 confocal microscope. Worms were grown in the presence of their respective ncAA at a concentration of 1 mM for 24 hours and illuminated with the 405 nm laser at 50% power. The worms were then imaged 24 hours after illumination. Scale bars represent: 20 µm.

**Supplementary Table 1 Transgenic *C. elegans* strains.**

| Name | Description | Genetic Back-ground | Expression Plasmids (µg) | Location in article |
|---|---|---|---|---|
| **SGR129** | greEx114[glr-1p::FLP(G5D)(K223TAG)::HA::egl-13NLS::SL2::mKate2; maco-1p::FRT_lacZ 3'UTR_FRT::Citrine2; sur-5p::HygR; glr-1p::smad-4NES::PCKRS(13IPYER); rpr-1p::tRNA(M15)] | N2 | KB438(6), KB454(2.5), JJ466(1.5), SE149(4.5) | Fig. 3 a, Fig. 5 a, b, Supplementary Fig. 4c, Supplementary Fig. 5 |
| **SGR130** | greEx115[glr-1p::FLP(G5D)(K223TAG)::HA::egl-13NLS::SL2::mKate2; maco-1p::FRT_lacZ 3'UTR_FRT::Citrine2; sur-5p::HygR; glr-1p::smad-4NES::PCKRS(13IPYER); rpr-1p::tRNA(M15)] | N2 | KB438(6), KB454(2.5), JJ466(1.5), SE149(4.5) | Fig. 3 a, Fig. 5 a, b, Supplementary Fig. 4c, Supplementary Fig. 5 |
| **SGR131** | greEx116[glr-1p::FLP(G5D)(K223TAG)::HA::egl-13NLS::SL2::mKate2; maco-1p::FRT_lacZ 3'UTR_FRT::Citrine2; sur-5p::HygR; glr-1p::smad-4NES::PCKRS(13IPYER); rpr-1p::tRNA(M15)] | N2 | KB438(6), KB454(2.5), JJ466(1.5), SE149(4.5) | Fig. 3 a, Fig. 5 a, b, Supplementary Fig. 4c, Supplementary Fig. 5 |
| **SGR132** | greEx117[glr-1p::FLP(G5D)(K223TAG)::HA::egl-13NLS::SL2::mKate2; maco-1p::FRT_lacZ 3'UTR_FRT::Citrine2; sur-5p::HygR; glr-1p::smad-4NES::PCKRS(13IPYER); rpr-1p::tRNA(M15)] | N2 | KB438(6), KB454(2.5), JJ466(1.5), SE149(4.5) | Fig. 2 c, Fig. 3 a, Fig. 5 a, b, Supplementary Fig. 3, Supplementary Fig. 4 c, Supplementary Fig. 5, Supplementary Fig. 9 d |
| **SGR112** | greEx97[glr-1p::FLP(G5D)(Y343TAG)::HA::egl-13NLS::SL2::mKate2; maco-1p::FRT_lacZ 3'UTR_FRT:::Citrine2; sur-5p::HygR; glr-1::Smad4::NPYRS; rpr-1p::tRNA(C15)] | N2 | KB463(6), KB454(2.5), AT11(1.5), SE150(4.5) | Fig. 2 c, Fig. 3 a, b, Fig. 5 a, b, c, Supplementary Fig. 1, Supplementary Fig. 3, Supplementary Fig. 5, Supplementary Fig. 9 c, d |

| | | | | |
|---|---|---|---|---|
| **SGR122** | greEx107[glr-1p::FLP(G5D)(Y343TAG)::HA::egl-13NLS::SL2::mKate2; maco-1p::FRT_lacZ 3'UTR_FRT::Citrine2; sur-5p::HygR; glr-1::Smad4::NPYRS; rpr-1p::tRNA(C15)] | N2 | KB463(6), KB454(2.5), AT11(1.5), SE150(4.5) | Fig. 3 c, b, Fig. 5 a, b, Supplementary Fig. 5, Supplementary Fig. 9 c |
| **SGR133** | greEx118[glr-1p::FLP(G5D)(C189TAG)::HA::egl-13NLS::SL2::mKate2; maco-1p::FRT_lacZ 3'UTR_FRT::Citrine2; sur-5p::HygR; glr-1p::smad-4NES::PCC2RS; rpr-1p::tRNA(M15)] | N2 | JJ543(6), KB454(2.5), JJ546(1.5), SE149(4.5) | Fig. 3 c, b, Fig. 5 a, b, Supplementary Fig. 5, Supplementary Fig. 9 c |
| **SGR134** | greEx119[glr-1p::FLP(G5D)(C189TAG)::HA::egl-13NLS::SL2::mKate2; maco-1p::FRT_lacZ 3'UTR_FRT::Citrine2; sur-5p::HygR; glr-1p::smad-4NES::PCC2RS; rpr-1p::tRNA(M15)] | N2 | JJ543(6), KB454(2.5), JJ546(1.5), SE149(4.5) | Fig. 3 c, b, Fig. 5 a, b, Supplementary Fig. 5, Supplementary Fig. 9 c |
| **SGR135** | greEx120[glr-1p::FLP(G5D)(C189TAG)::HA::egl-13NLS::SL2::mKate2; maco-1p::FRT_lacZ 3'UTR_FRT::Citrine2; sur-5p::HygR; glr-1p::smad-4NES::PCC2RS; rpr-1p::tRNA(M15)] | N2 | JJ543(6), KB454(2.5), JJ546(1.5), SE149(4.5) | Fig. 2 c, Fig. 3 a, Fig. 5 a, b, Supplementary Fig. 3, Supplementary Fig. 5, Supplementary Fig. 9 c, d |
| **SGR136** | greEx121[glr-1p::FLP(G5D)(C189TAG)::HA::egl-13NLS::SL2::mKate2; maco-1p::FRT_lacZ 3'UTR_FRT::Citrine2; sur-5p::HygR; glr-1p::smad-4NES::PCC2RS; rpr-1p::tRNA(M15)] | N2 | JJ543(6), KB454(2.5), JJ546(1.5), SE149(4.5) | Fig. 3 a, Fig. 5 a, b, Supplementary Fig. 5, Supplementary Fig. 9 c |
| **SGR137** | greEx122[glr-1p::FLP(G5D)(C189TAG)::HA::egl-13NLS::SL2::mKate2; maco-1p::FRT_lacZ 3'UTR_FRT::Citrine2; sur-5p::HygR; glr-1p::smad-4NES::PCC2RS; rpr-1p::tRNA(M15)] | N2 | JJ543(6), KB454(2.5), JJ546(1.5), SE149(4.5) | Fig. 3 a, Fig. 5 a, b, Supplementary Fig. 5, Supplementary Fig. 9 c |
| **SGR138** | greEx123[glr-1p::FLP(G5D)(K223TAG)::HA::egl-13NLS::SL2::mKate2; maco-1p::FRT_lacZ 3'UTR_FRT::Citrine2; sur-5p::HygR; glr-1p::smad-4NES::HCKRS; rpr-1p::tRNA(M15)] | N2 | KB438(6), KB454(2.5), JJ461(1.5), SE149(4.5) | Supplementary Fig. 4 c |
| **SGR139** | greEx124[glr-1p::FLP(G5D)(K223TAG)::HA::egl-13NLS::SL2::mKate2; maco-1p::FRT_lacZ 3'UTR_FRT::Citrine2; sur-5p::HygR; glr-1p::smad-4NES::HCKRS; rpr-1p::tRNA(M15)] | N2 | KB438(6), KB454(2.5), JJ461(1.5), SE149(4.5) | Supplementary Fig. 4 c |
| **SGR140** | greEx125[glr-1p::FLP(G5D)(K223TAG)::HA::egl-13NLS::SL2::mKate2; maco-1p::FRT_lacZ 3'UTR_FRT::Citrine2; sur-5p::HygR; glr-1p::smad-4NES::HCKRS; rpr-1p::tRNA(M15)] | N2 | KB438(6), KB454(2.5), JJ461(1.5), SE149(4.5) | Supplementary Fig. 4 c |
| **SGR141** | greEx126[glr-1p::Cre(R119A)(Y324TAG)::HA::egl-13NLS::SL2::mKate2; maco-1p::loxP_lacZ | N2 | JJ191(6), KB462(2.5), | Fig. 3 a, b, Supplementary Fig. 5, |

| | | | | |
|---|---|---|---|---|
| | 3'UTR_loxP::Citrine2; sur-5p::HygR; glr-1p::smad-4NES::NPYRS; rpr-1p::tRNA(M15)] | | AT11(1.5), SE149(4.5) | Supplemen-tary Fig. 1 |
| **SGR142** | greEx127[glr-1p::Cre(R119A)(Y324TAG)::HA::egl-13NLS::SL2::mKate2; maco-1p::loxP_lacZ 3'UTR_loxP::Citrine2; sur-5p::HygR; glr-1p::smad-4NES::NPYRS; rpr-1p::tRNA(M15)] | N2 | JJ191(6), KB462(2.5), AT11(1.5), SE149(4.5) | Fig. 3 a, b, Supplemen-tary Fig. 5 |
| **SGR143** | greEx128[glr-1p::Cre(R119A)(K201TAG)::HA::egl-13NLS::SL2::mKate2; maco-1p::loxP_lacZ 3'UTR_loxP::Citrine2; sur-5p::HygR; glr-1p::smad-4NES::PCKRS(13IPYER); rpr-1p::tRNA(M15)] | N2 | KB389(6), KB462(2.5), JJ466(1.5), SE149(4.5) | Fig. 3 a, Supplemen-tary Fig. 5 |
| **SGR144** | greEx129[glr-1p::Cre(R119A)(K201TAG)::HA::egl-13NLS::SL2::mKate2; maco-1p::loxP_lacZ 3'UTR_loxP::Citrine2; sur-5p::HygR; glr-1p::smad-4NES::PCKRS(13IPYER); rpr-1p::tRNA(M15)] | N2 | KB389(6), KB462(2.5), JJ466(1.5), SE149(4.5) | Fig. 3 a, Supplemen-tary Fig. 5 |
| **SGR107** | greEx92[glr-1p::FLP(G5D)(K223TAG)::HA::egl-13NLS::SL2::mKate2; maco-1p::FRT_lacZ 3'UTR_FRT::Chrimson::F2A::Citrine2; rps-0p::HygR; glr-1::Smad4::PCKRS; rpr-1p::tRNA(C15)] | N2 | KB438(6), KB443(2.5), SE174(1.5), SE150(4.5) | Fig. 3 c |
| **SGR118** | greEx103[glr-1p::FLP(G5D)(K223)TAG::HA::egl-13NLS::SL2::mKate2; maco-1p::FRT_lacZ 3'UTR_FRT::Chrimson::F2A::Citrine2; rps-0p::HygR; glr-1::Smad4::PCKRS; rpr-1p::tRNA(C15)] | N2 | KB438(6), KB443(2.5), SE174(1.5), SE150(4.5) | Fig. 3 c |
| **SGR149** | greEx134[glr-1p::FLP(G5D)(Y343TAG)::HA::egl-13NLS::SL2::mKate2; maco-1p::FRT_lacZ 3'UTR_FRT::Chrimson::F2A::Citrine2; sur-5p::HygR; glr-1p::smad-4NES::NPYRS; rpr-1p::tRNA(M15)] | N2 | KB463(6), KB458(2.5), AT11(1.5), SE149(4.5) | Fig. 3 c |
| **SGR150** | greEx135[glr-1p::FLP(G5D)(C189TAG)::HA::egl-13NLS::SL2::mKate2; maco-1p::FRT_lacZ 3'UTR_FRT::Chrimson::F2A::Citrine2; sur-5p::HygR; glr-1p::smad-4NES::PCC2RS; rpr-1p::tRNA(M15)] | N2 | JJ546(6), KB458(2.5), JJ546(1.5), SE149(4.5) | Fig. 3 c |
| **SGR151** | greEx136[glr-1p::FLP(G5D)(C189TAG)::HA::egl-13NLS::SL2::mKate2; maco-1p::FRT_lacZ 3'UTR_FRT::Chrimson::F2A::Citrine2; sur-5p::HygR; glr-1p::smad-4NES::PCC2RS; rpr-1p::tRNA(M15)] | N2 | JJ546(6), KB458(2.5), JJ546(1.5), SE149(4.5) | Fig. 3 c |
| **SGR152** | greEx137[glr-1p::FLP(G5D)(C189TAG)::HA::egl-13NLS::SL2::mKate2; maco-1p::FRT_lacZ 3'UTR_FRT::Chrimson::F2A::Citrine2; sur-5p::HygR; glr-1p::smad-4NES::PCC2RS; rpr-1p::tRNA(M15)] | N2 | JJ546(6), KB458(2.5), JJ546(1.5), SE149(4.5) | Fig. 3 c |
| **SGR197** | greEx178[glr-1p::FLP(G5D)(K303TAG)::HA::egl- | N2 | JJ466(1.5), JJ544(6) | Methods - Imaging and analysis |

| | | | | |
|---|---|---|---|---|
| | 13NLS::SL2::mKate2; sur-5p::HygR; glr-1p::smad-4NES::PCKRS(13IPYER)] | | | |
| **SGR153** | greEx138[sur-5p::smad-4NES::PCKRS(wt); sur-5p::HygR; rps--0p::GFP(TAG)::mCherry::HA::egl-13NLS; rpr-1p::tRNA(M15)] | smg-6(ok1794) | JJ518(2), SG88(3), SE149(6) | Supplementary Fig. 4 b |
| **SGR154** | greEx139[sur-5p::smad-4NES::PCKRS(wt); sur-5p::HygR; rps--0p::GFP(TAG)::mCherry::HA::egl-13NLS; rpr-1p::tRNA(M15)] | smg-6(ok1794) | JJ518(2), SG88(3), SE149(6) | Supplementary Fig. 4 b |
| **SGR155** | greEx140[sur-5p::smad-4NES::PCKRS(13IPYER); sur-5p::HygR; rps-0p::GFP(TAG)::mCherry::HA::egl-13NLS; rpr-1p::tRNA(M15)] | smg-6(ok1794) | JJ238(2), SG88(3), SE149(7) | Supplementary Fig. 4 b |
| **SGR156** | greEx141[sur-5p::smad-4NES::PCKRS(13IPYER); sur-5p::HygR; rps-0p::GFP(TAG)::mCherry::HA::egl-13NLS; rpr-1p::tRNA(M15)] | smg-6(ok1794) | JJ238(2), SG88(3), SE149(7) | Supplementary Fig. 4 b |
| **SGR157** | greEx142[sur-5p::smad-4NES::PCKRS(13IPYER); sur-5p::HygR; rps-0p::GFP(TAG)::mCherry::HA::egl-13NLS; rpr-1p::tRNA(M15)] | smg-6(ok1794) | JJ238(2), SG88(3), SE149(7) | Supplementary Fig. 4 b |
| **SGR158** | greEx143[sur-5p::smad-4NES::HCKRS; sur-5p::HygR; rps--0p::GFP(TAG)::mCherry::HA::egl-13NLS; rpr-1p::tRNA(M15)] | smg-6(ok1794) | JJ345(2), SG88(3), SE149(7) | Supplementary Fig. 4 b |
| **SGR159** | greEx144[sur-5p::smad-4NES::HCKRS; sur-5p::HygR; rps--0p::GFP(TAG)::mCherry::HA::egl-13NLS; rpr-1p::tRNA(M15)] | smg-6(ok1794) | JJ345(2), SG88(3), SE149(7) | Supplementary Fig. 4 b |
| **SGR160** | greEx145[sur-5p::smad-4NES::HCKRS; sur-5p::HygR; rps--0p::GFP(TAG)::mCherry::HA::egl-13NLS; rpr-1p::tRNA(M15)] | smg-6(ok1794) | JJ345(2), SG88(3), SE149(7) | Supplementary Fig. 4 b |
| **SGR203** | greEx184[glr-1p::FLP(G5D)(Y343TAG)::HA::egl-13NLS::SL2::mTagBFP2; sur-5p::HygR; glr-1p::smad-4NES::NPYRS; rpr-1p::tRNA(M15)] | BN294 | JJ740(6), AT11(1.5), SE149(4.5) | Fig. 3 d, Supplementary Fig. 7 b |
| **SGR204** | greEx185[glr-1p::FLP(G5D)(Y343TAG)::HA::EGL-13_NLS::SL2:EGL-13_NLS:::mTagBFP2; sur-5p::HygR; glr-1p::smad-4NES::NPYRS; rpr-1p::tRNA(M15)] | BN294 | JJ740(6), AT11(1.5), SE149(4.5) | Fig. 3 d, Supplementary Fig. 7 b |
| **SGR205** | greEx186[myo-3p::PKIa::MmNPYRS; sur-5p::HygR;; rpr-1P::tRNA(C15); myo-3p::FLP(G5D)(Y343TAG) egl-13 SL2 mKate2 let-858; eft-3p::FRT_bGal_FRT Citrine2_HA let-858] | N2 | KB329(1.5), SE149(4.5), JJ660(6), KB450(2.5) | Fig. 4 a, Supplementary Fig. 8 a |
| **SGR206** | greEx187[myo-2p::PKIa::MmNPYRS; sur-5p::HygR;; rpr-1P::tRNA(C15); myo-2p::FLP(G5D)(Y343TAG) egl-13 SL2 mKate2 let-858; eft-3p::FRT_bGal_FRT Citrine2_HA let-858] | N2 | JJ661(1.5), SE149(4.5), JJ659(6), KB450(2.5) | Fig. 4 b, Supplementary Fig. 8 b |
| **SGR207** | greEx188[myo-3p::PKIa::MmNPYRS; sur-5p::HygR;; rpr-1P::tRNA(C15); myo-3p::Cre_(Y324TAG)egl-13 SL2 mKate2 let-858; eft-3p::loxP_bGal_loxP Citrine2_HA let-858] | N2 | KB329(1.5), SE149(4.5), JJ658(6), JJ656(2.5) | Fig. 4. a, Supplementary Fig. 8 a |

| | | | | |
|---|---|---|---|---|
| **SGR208** | greEx189[myo-2p::PKIa::MmNPYRS; sur-5p::HygR;; rpr-1P::tRNA(C15); myo-2p::Cre_(Y324TAG)egl-13 SL2 mKate2 let-858; eft-3p::loxP_bGal_loxP Citrine2_HA let-858] | N2 | JJ661(1.5), SE149(4.5), JJ657(6), JJ656(2.5) | Fig. 4 b, Supplementary Fig. 8 b |

**Supplementary Table 2 Expression plasmids.**

| Plasmid Number | Description | Source Plasmids | | | |
|---|---|---|---|---|---|
| | | Dest | p1 | p2 | p3 |
| **AT11** | *glr-1p::HuSmad4::MmNPYRS::let858_3'UTR; sur-5p::HygR::unc-54_3'UTR* | SE80 | IR157 | KB126 | SG606 |
| **JJ191** | *glr-1p::Cre(R119A,Y324TAG)::HA::EGL-13_NLS::SL2::mKate2::let858_3'UTR* | PDESTII | IR157 | JJ190 | IR322 |
| **JJ238** | *sur-5p::HuSmad4::MmP-CKRS(13IPYER)::let858_3'UTR; sur-5p::HygR::unc-54_3'UTR* | SE80 | SE72 | JJ237 | SG606 |
| **JJ345** | *sur-5p::HuSmad4::MmHCKRS::let858_3'UTR; sur-5p::HygR::unc-54_3'UTR* | SE80 | SE72 | JJ338 | SG606 |
| **JJ461** | *glr-1p::HuSmad4::MmHCKRS::let858_3'UTR; sur-5p::HygR::unc-54_3'UTR* | SE80 | IR157 | JJ338 | SG606 |
| **JJ466** | *glr-1p::HuSmad4::MmP-CKRS(13IPYER)::let858_3'UTR; sur-5p::HygR::unc-54_3'UTR* | SE80 | IR157 | JJ237 | SG606 |
| **JJ518** | *sur-5p::HuSmad4::PCKRS::let858_3'UTR; sur-5p::HygR::unc-54_3'UTR* | SE80 | SE72 | SE154 | SG606 |
| **JJ543** | *glr-1p::FLP(G5D,C189TAG)::HA::EGL-13_NLS::SL2::mKate2::let858_3'UTR* | PDESTII | IR157 | JJ540 | IR322 |
| **JJ544** | *glr-1p::FLP(G5D,K303TAG)::HA::EGL-13_NLS::SL2::mKate2::let858_3'UTR* | PDESTII | IR157 | JJ541 | IR322 |
| **JJ546** | *glr-1p::HuSmad4::MmPCC2RS::let858_3'UTR; sur-5p::HygR::unc-54_3'UTR* | SE80 | IR157 | KB124 | SG606 |
| **JJ740** | *glr-1p::FLP(G5D)(Y343TAG)::HA::EGL-13_NLS::SL2:EGL-13NLS:::mTagBFP2* | PDESTII | IR157 | KB321 | JOS163 |
| **KB389** | *glr-1p::Cre(R119A,K201TAG)::HA::EGL-13_NLS::SL2::mKate2::let858_3'UTR* | PDESTII | IR157 | SE17 | IR322 |
| **KB438** | *glr-1p::FLP(G5D,K223TAG)::HA::EGL-13_NLS::SL2::mKate2::let858_3'UTR* | PDESTII | IR157 | KB257 | IR322 |

| | | | | | |
|---|---|---|---|---|---|
| **KB443** | *maco-1p::FRT::B-Gal_3'UTR::FRT::Chrimson::F2A::Citrine2::HA::let858_3'UTR* | PDESTII | SE77 | LD222 | KB431 |
| **KB454** | *maco-1p::FRT::B-Gal_3'UTR::FRT::Citrine2::HA::let858_3'UTR* | PDESTII | SE77 | LD222 | KB442 |
| **KB458** | *maco-1p::FRT::B-Gal_3'UTR::FRT::Chrimson::F2A::Citrine2::HA::egl-13::let858_3'UTR* | PDESTII | SE77 | LD222 | KB432 |
| **KB462** | *maco-1p::loxp::B-Gal-3'UTR::loxp::Citrine2::HA::let858_3'UTR* | PDESTII | SE77 | SG365 | KB442 |
| **KB463** | *glr-1p::FLP(G5D,Y343TAG)::HA::EGL-13_NLS::SL2::mKate2::let858_3'UTR* | PDESTII | IR157 | KB321 | IR322 |
| **SE149** | *rpr-1p::M15(CUA)::sup-7_3'UTR* | [3] | | | |
| **SE150** | *rpr-1p::C15(CUA)::sup-7_3'UTR* | [4] | | | |
| **SE174** | *glr-1p::HuSmad4::MmP-CKRS::let858_3'UTR; sur-5p::HygR::unc-54_3'UTR* | [4] | | | |
| **SG88** | *rps-0p::GFP::TAG::mCherry::HA::EGL-13_NLS::unc-54_3'UTR* | [5] | | | |

**Supplementary Table 3 Gateway Entry plasmids.**

| Plasmid Number | Description | Source |
|---|---|---|
| **IR157** | *glr-1p* | [4] |
| **IR322** | *SL2::mKate2::let858_3'UTR* | IR322 is identical to IR182[4] with an mKate2 gene in place of the GFP gene |
| **JJ190** | *Cre(R119A,Y324TAG)::HA::EGL-13_NLS* | Modified from SE17[4], K201TAG removed, and Y324TAG introduced with primers [(F) aacatcgtcatgaactagatccgtaacctcgactc, (R) ctagttcatgacgatgttgacgttg] |
| **JJ237** | *HuSmad4::MmPCKRS(13IPYER)* | Constructed by introducing mutations T13I, T56P, H62Y, A100E, S193R into SE154 |
| **JJ338** | *HuSmad4::MmHCKRS* | Constructed by introducing mutations Y306A and L309M into HuSmad4::MmPylRS using primers [(F) CTCGCCAACTACATGCGTAA-GCTCGACCGTGCC, (R) CATGTAGTT-GGCGAGGTTTGGGGCGAGCATTG] |
| **JJ540** | *FLP(G5D,C189TAG)::HA::EGL-13_NLS* | Constructed by introducing C189TAG into KB258 FLP(G5D)::HA::egl-13 using primers [(F) catcaacTAGggacgtttctccgacatcaagaac, (R) gagaaacgtccCTAgttgatgaaggtggcgag-gaag] |
| **JJ541** | *FLP(G5D,K303TAG)::HA::EGL-13_NLS* | Constructed by introducing K303 into KB258 FLP(G5D)::HA::egl-13 using primers [(Fgaa-cggaccaTagtcccacatcggacgtcac, (R) gatgtgg-gactAtggtccgttcttgatggcgaag] |
| **JOS163** | *SL2::EGL-13NLS:::mTagBFP2* | [6] |
| **KB124** | *HuSmad4::MmPCC2RS* | [3] |
| **KB126** | *HuSmad4::MmNPYRS* | Identical to KB125[6], but with HuSmad4 instead of Pkiα |
| **KB257** | *FLP(G5D,K223TAG)::HA::EGL-13_NLS* | Constructed by introducing G5D mutation into SE13 FLP(K223TAG)::HA::egl-13 using primers [(F) |

| | | |
|---|---|---|
| | | acaattcgACatcctctgcaagaccccaccaaag, (R) cttgcagaggatGTcgaattgtggcatttttagcctg] |
| **KB321** | *FLP(G5D,Y343TAG)::HA::EGL-13_NLS* | Constructed by introducing Y343TAG into KB258 FLP(G5D)::HA::egl-13 using primers [(F) gtaccaccTAGacccaccaaatcaccgccatcccagac, (R) gatggcggtgatttggtgggtCTAggtggtacgggcgacggcggaggcacgcttg] |
| **KB431** | *Chrimson::F2A::Citrine2::HA::let858_3'UTR* | Constructed by Nebuilder between amplification of Chrimson::Citrine2::let-858_3’UTR using primers [(F) GTCGAGTCCAACCCAGGACCAATGGTATCGAAAGGaGAGG, (R) TGCTTGACTCCGGATCCTGGGACTGTATCCTCATCTTCCTC] and oligonucleotide [TGGTCCTGGGTTGGACTCGACGTCTCCGGCGAGCTTGAGGAGGTCGAAGTTGAGGGTTTGCTTGACTCCGGATCCTGG] to introduce the F2A |
| **KB432** | *Chrimson::F2A::Citrine2::HA::egl-13::let858_3'UTR* | Constructed by introducing egl-13 into KB431 using primers [(F) TCCGACAAAACTGAGTGAAAACGCGAAGAAGCTTGCCAAGGAAGTTGAAAATtaacgtgaagtggaatcggatgatcgacgccg, (R) GCTTCTTCGCGTTTTCACTCAGTTTTGTCGGATTCGCTTTTCGTCTACGGCTCATAGCGTAGTCTGGGACATCATATGGGTA] |
| **KB442** | *Citrine2::HA::let858_3'UTR* | Constructed by BP reaction into pDONR P2r-P3 of fragment generated by PCR from Chrimson::Citrine2::let-858_3’UTR using primers [(F) GGGGACAGCTTTCTTGTACAAAGTGGTAAAAAAATGGTATCGAAAGGAGAGGAATTATTTACTG, (R) GGGGACAACTTTGTATAATAAAGTTGatacggattcgcatttgccaag] |
| **LD222** | *FRT::B-Gal_3'UTR::FRT* | Constructed by BP reaction into pDONR221 of fragment generated by PCR from SG365 loxp::B-Gal-3'UTR::loxp using primers [(F) GGGGACAAGTTTGTACAAAAAAGCAGGCTgaagttcctatactttctagagaataggaacttcggaataggaacttcATGGTCGTTTTAC AACGTCGTGACTGGGAAAACCC, (R) GGGGACCACTTTGTACAAGAAAGCTGGGTgaagttcctattccgaagttcctattctctagaaagtataggaacttcGTTTCATGTTTCAAAAAA AATTTATTCTTTATTCTTTATTAGATCCTATTATTTTTGACACCAGACAAGTTGGTAATGGTAGC] using SG365 (Davis2021) as a template |
| **SE154** | *HuSmad4::PCKRS* | [4] |
| **SE17** | *Cre(R119A,K201TAG)* | [4] |
| **SE72** | *sur-5p* | [4] |
| **SE77** | *maco-1p* | [4] |
| **SE80** | *sur-5p::HygR::unc-54_3'UTR* | Constructed from IR98[4] by replacing the rps-0p::HygR::unc-54_3’UTR with sur-5p::HygR::unc-54_3’UTR by ligation using restriction enzymes KasI and AscI |
| **SG365** | *loxp::B-Gal-3'UTR::loxp* | [4] |

| | | |
|---|---|---|
| **SG606** | *let858_3'UTR* | 4 |

**Supplementary Note 1 Gene sequences.**

Amber (TAG) stop codons to direct ncAA incorporation are marked in red, bolded and underlined. Mutations in PCKRS(13IPYER) as compared to PCKRS are also marked in red, bolded and underlined. Artificial introns are marked in green and lowercase. The nuclear localization sequence (NLS) from C. elegans EGL-13 is marked in red. The HA tag is marked in blue. The nuclear export sequence from human Smad4 is marked in purple and underlined.

**FLP(Y343TAG)::HA::ELG-13_NLS**
ATGCCACAATTCGACATCCTCTGCAAGACCCCACCAAAGGTCCTCGTCCGTCAATTCGTCGAGCG
TTTCGAGCGTCCATCCGGAGAGAAGATCGCCCTCTGCGCCGCCGAGCTCACCTACCTCTGCTGG
ATGATCACCCACAACGGAACCGCCATCAAGCGTGCCACCTTCATGTCCTACAACACCATCATCTC
CAACTCCCTCTCCTTCGACATCGTCAACAAGTCCCTCCAATTCAAGTACAAGACCCAAAAGGCCA
CCATCCTCGAGGCCTCCCTCAAGAAGCTCATCCCAGCCTGGGAGTTCACCATCATCCCATACTAC
GGACAAAAGCACCAATCCGACATCACCGACATCGTCTCCTCCCTCCAACTCCAATTCGAGTCCTC
CGAGGAGGCCGACAAGgtaagtttaaacatatatatactaactaaccctgattatttaaattttcagGGAAACTCCCACTCC
AAGAAGATGCTCAAGGCCCTCCTCTCCGAGGGAGAGTCCATCTGGGAGATCACCGAGAAGATCC
TCAACTCCTTCGAGTACACCTCCCGTTTCACCAAGACCAAGACCCTCTACCAATTCCTCTTCCTCG
CCACCTTCATCAACTGCGGACGTTTCTCCGACATCAAGAACGTCGACCCAAAGTCCTTCAAGCTC
GTCCAAAACAAGTACCTCGGAGTCATCATCCAATGCCTCGTCACCGAGACCAAGACCTCCGTCTC
CCGTCACATCTACTTCTTCTCCGCCCGTGGACGTATCGACCCACTCGTCTACCTCGACGAGTTCC
TCCGTAACTCCGAGCCAGTCCTCAAGCGTGTCAACCGTACCGGAAACTCCTCCTCCAACAAGCA
AGAGTACCAACTCCTCAAGGACAACCTCGTCCGTTCCTACAACAAGgtaagtttaaacagttcggtactaacta
accatacatatttaaattttcagGCCCTCAAGAAGAACGCCCCATACTCCATCTTCGCCATCAAGAACGGACC
AAAGTCCCACATCGGACGTCACCTCATGACCTCCTTCCTCTCCATGAAGGGACTCACCGAGCTCA
CCAACGTCGTCGGAAACTGGTCCGACAAGCGTGCCTCCGCCGTCGCCCGTACCACC**TAG**ACCC
ACCAAATCACCGCCATCCCAGACCACTACTTCGCCCTCGTCTCCCGTTACTACGCCTACGACCCA
ATCTCCAAGGAGATGATCGCCCTCAAGGACGAGACCAACCCAATCGAGGAGTGGCAACACATCG
AGCAACTCAAGGGATCCGCCGAGGGATCCATCCGTTACCCAGCCTGGAACGGAATCATCTCCCA
AGAGGTCCTCGACTACCTCTCCTCCTACATCAACCGTCGTATCTACCCATATGATGTCCCAGACTA
CGCTATGAGCCGTAGACGAAAAGCGAATCCGACAAAACTGAGTGAAAACGCGAAGAAGCTTGCC
AAGGAAGTTGAAAATTAA

HA
EGL-13 NLS

**FLP(K223TAG)::HA::ELG-13_NLS**
ATGCCACAATTCGACATCCTCTGCAAGACCCCACCAAAGGTCCTCGTCCGTCAATTCGTCGAGCG
TTTCGAGCGTCCATCCGGAGAGAAGATCGCCCTCTGCGCCGCCGAGCTCACCTACCTCTGCTGG
ATGATCACCCACAACGGAACCGCCATCAAGCGTGCCACCTTCATGTCCTACAACACCATCATCTC
CAACTCCCTCTCCTTCGACATCGTCAACAAGTCCCTCCAATTCAAGTACAAGACCCAAAAGGCCA
CCATCCTCGAGGCCTCCCTCAAGAAGCTCATCCCAGCCTGGGAGTTCACCATCATCCCATACTAC
GGACAAAAGCACCAATCCGACATCACCGACATCGTCTCCTCCCTCCAACTCCAATTCGAGTCCTC
CGAGGAGGCCGACAAGgtaagtttaaacatatatatactaactaaccctgattatttaaattttcagGGAAACTCCCACTCC
AAGAAGATGCTCAAGGCCCTCCTCTCCGAGGGAGAGTCCATCTGGGAGATCACCGAGAAGATCC
TCAACTCCTTCGAGTACACCTCCCGTTTCACCAAGACCAAGACCCTCTACCAATTCCTCTTCCTCG
CCACCTTCATCAACTGCGGACGTTTCTCCGACATCAAGAACGTCGACCCAAAGTCCTTCAAGCTC
GTCCAAAACAAGTACCTCGGAGTCATCATCCAATGCCTCGTCACCGAGACC**TAG**ACCTCCGTCTC
CCGTCACATCTACTTCTTCTCCGCCCGTGGACGTATCGACCCACTCGTCTACCTCGACGAGTTCC
TCCGTAACTCCGAGCCAGTCCTCAAGCGTGTCAACCGTACCGGAAACTCCTCCTCCAACAAGCA
AGAGTACCAACTCCTCAAGGACAACCTCGTCCGTTCCTACAACAAGgtaagtttaaacagttcggtactaacta
accatacatatttaaattttcagGCCCTCAAGAAGAACGCCCCATACTCCATCTTCGCCATCAAGAACGGACC
AAAGTCCCACATCGGACGTCACCTCATGACCTCCTTCCTCTCCATGAAGGGACTCACCGAGCTCA
CCAACGTCGTCGGAAACTGGTCCGACAAGCGTGCCTCCGCCGTCGCCCGTACCACCTACACCC
ACCAAATCACCGCCATCCCAGACCACTACTTCGCCCTCGTCTCCCGTTACTACGCCTACGACCCA
ATCTCCAAGGAGATGATCGCCCTCAAGGACGAGACCAACCCAATCGAGGAGTGGCAACACATCG

AGCAACTCAAGGGATCCGCCGAGGGATCCATCCGTTACCCAGCCTGGAACGGAATCATCTCCCA
AGAGGTCCTCGACTACCTCTCCTCCTACATCAACCGTCGTATCTACCCATATGATGTCCCAGACTA
CGCTATGAGCCGTAGACGAAAAGCGAATCCGACAAAACTGAGTGAAAACGCGAAGAAGCTTGCC
AAGGAAGTTGAAAATTAA

HA
EGL-13 NLS

**FLP(C189TAG)::HA::ELG-13_NLS**
ATGCCACAATTCGACATCCTCTGCAAGACCCCACCAAAGGTCCTCGTCCGTCAATTCGTCGAGCG
TTTCGAGCGTCCATCCGGAGAGAAGATCGCCCTCTGCGCCGCCGAGCTCACCTACCTCTGCTGG
ATGATCACCCACAACGGAACCGCCATCAAGCGTGCCACCTTCATGTCCTACAACACCATCATCTC
CAACTCCCTCTCCTTCGACATCGTCAACAAGTCCCTCCAATTCAAGTACAAGACCCAAAAGGCCA
CCATCCTCGAGGCCTCCCTCAAGAAGCTCATCCCAGCCTGGGAGTTCACCATCATCCCATACTAC
GGACAAAAGCACCAATCCGACATCACCGACATCGTCTCCTCCCTCCAACTCCAATTCGAGTCCTC
CGAGGAGGCCGACAAGgtaagtttaaacatatatatactaactaaccctgattatttaaattttcagGGAAACTCCCACTCC
AAGAAGATGCTCAAGGCCCTCCTCTCCGAGGGAGAGTCCATCTGGGAGATCACCGAGAAGATCC
TCAACTCCTTCGAGTACACCTCCCGTTTCACCAAGACCAAGACCCTCTACCAATTCCTCTTCCTCG
CCACCTTCATCAAC**TAG**GGACGTTTCTCCGACATCAAGAACGTCGACCCAAAGTCCTTCAAGCTC
GTCCAAAACAAGTACCTCGGAGTCATCATCCAATGCCTCGTCACCGAGACCAAGACCTCCGTCTC
CCGTCACATCTACTTCTTCTCCGCCCGTGGACGTATCGACCCACTCGTCTACCTCGACGAGTTCC
TCCGTAACTCCGAGCCAGTCCTCAAGCGTGTCAACCGTACCGGAAACTCCTCCTCCAACAAGCA
AGAGTACCAACTCCTCAAGGACAACCTCGTCCGTTCCTACAACAAGgtaagtttaaacagttcggtactaacta
accatacatatttaaattttcagGCCCTCAAGAAGAACGCCCCATACTCCATCTTCGCCATCAAGAACGGACC
AAAGTCCCACATCGGACGTCACCTCATGACCTCCTTCCTCTCCATGAAGGGACTCACCGAGCTCA
CCAACGTCGTCGGAAACTGGTCCGACAAGCGTGCCTCCGCCGTCGCCCGTACCACCTACACCC
ACCAAATCACCGCCATCCCAGACCACTACTTCGCCCTCGTCTCCCGTTACTACGCCTACGACCCA
ATCTCCAAGGAGATGATCGCCCTCAAGGACGAGACCAACCCAATCGAGGAGTGGCAACACATCG
AGCAACTCAAGGGATCCGCCGAGGGATCCATCCGTTACCCAGCCTGGAACGGAATCATCTCCCA
AGAGGTCCTCGACTACCTCTCCTCCTACATCAACCGTCGTATCTACCCATATGATGTCCCAGACTA
CGCTATGAGCCGTAGACGAAAAGCGAATCCGACAAAACTGAGTGAAAACGCGAAGAAGCTTGCC
AAGGAAGTTGAAAATTAA

HA
EGL-13 NLS

**Cre(Y324TAG)::HA::ELG-13_NLS**
ATGTCGAATTTGCTCACCGTCCACCAAAACCTCCCAGCCCTCCCAGTCGACGCCACCTCCGACG
AGGTCCGTAAGAACCTCATGGACATGTTCCGTGACCGTCAAGCCTTCTCCGAGCACACCTGGAA
GATGCTCCTCTCCGTCTGCCGTTCCTGGGCCGCCTGGTGCAAGCTCAACAACCGTAAGTGGTTC
CCAGCCGAGCCAGAGGACGTCCGTGACTACCTCCTCTACCTCCAAGCCCGTGGACTCGCCGTCA
AGgtaagtttaaacatatatatactaactaaccctgattatttaaattttcagACCATCCAACAACACCTCGGACAACTCAAC
ATGCTCCACCGTCGTTCCGGACTCCCACGTCCATCCGACTCCAACGCCGTCTCCCTCGTCATGC
GTGCTATCCGTAAGGAGAACGTCGACGCCGGAGAGCGTGCCAAGCAAGCCCTCGCCTTCGAGC
GTACCGACTTCGACCAAGTCCGTTCCCTCATGGAGAACTCCGACCGTTGCCAAGACATCCGTAAC
CTCGCCTTCCTCGGAATCGCCTACAACACCCTCCTCCGTATCGCCGAGATCGCCCGTATCCGTGT
CAAGgtaagtttaaacagttcggtactaactaaccatacatatttaaattttcagGACATCTCCCGTACCGACGGAGGACGT
ATGCTCATCCACATCGGACGTACCAAGACCCTCGTCTCCACCGCCGGAGTCGAGAAGGCCCTCT
CCCTCGGAGTCACCAAGCTCGTCGAGCGTTGGATCTCCGTCTCCGGAGTCGCCGACGACCCAA
ACAACTACCTCTTCTGCCGTGTCCGTAAGgtaagtttaaacatgattttactaactaactaatctgatttaaattttcagAACG
GAGTCGCCGCCCCATCCGCCACCTCCCAACTCTCCACCCGTGCCCTCGAGGGAATCTTCGAGG
CCACCCACCGTCTCATCTACGGAGCCAAGGACGACTCCGGACAACGTTACCTCGCCTGGTCCGG
ACACTCCGCCCGTGTCGGAGCCGCCCGTGACATGGCCCGTGCCGGAGTCTCCATCCCAGAGAT
CATGCAAGCCGGAGGATGGACCAACGTCAACATCGTCATGAAC**TAG**ATCCGTAACCTCGACTCCG
AGACCGGAGCCATGGTCCGTCTCCTCGAGGACGGAGACTACCCATATGATGTCCCAGACTACGC
TATGAGCCGTAGACGAAAAGCGAATCCGACAAAACTGAGTGAAAACGCGAAGAAGCTTGCCAAG
GAAGTTGAAAATTAA

HA
EGL-13 NLS

**Cre(K201TAG)::HA::ELG-13_NLS**
ATGTCGAATTTGCTCACCGTCCACCAAAACCTCCCAGCCCTCCCAGTCGACGCCACCTCCGACG
AGGTCCGTAAGAACCTCATGGACATGTTCCGTGACCGTCAAGCCTTCTCCGAGCACACCTGGAA
GATGCTCCTCTCCGTCTGCCGTTCCTGGGCCGCCTGGTGCAAGCTCAACAACCGTAAGTGGTTC
CCAGCCGAGCCAGAGGACGTCCGTGACTACCTCCTCTACCTCCAAGCCCGTGGACTCGCCGTCA
AGgtaagtttaaacatatatatactaactaaccctgattatttaaattttcagACCATCCAACAACACCTCGGACAACTCAAC
ATGCTCCACCGTCGTTCCGGACTCCCACGTCCATCCGACTCCAACGCCGTCTCCCTCGTCATGC
GTGCTATCCGTAAGGAGAACGTCGACGCCGGAGAGCGTGCCAAGCAAGCCCTCGCCTTCGAGC
GTACCGACTTCGACCAAGTCCGTTCCCTCATGGAGAACTCCGACCGTTGCCAAGACATCCGTAAC
CTCGCCTTCCTCGGAATCGCCTACAACACCCTCCTCCGTATCGCCGAGATCGCCCGTATCCGTGT
CAAGgtaagtttaaacagttcggtactaactaaccatacatatttaaattttcagGACATCTCCCGTACCGACGGAGGACGT
ATGCTCATCCACATCGGACGTACC**TAG**ACCCTCGTCTCCACCGCCGGAGTCGAGAAGGCCCTCT
CCCTCGGAGTCACCAAGCTCGTCGAGCGTTGGATCTCCGTCTCCGGAGTCGCCGACGACCCAA
ACAACTACCTCTTCTGCCGTGTCCGTAAGgtaagtttaaacatgattttactaactaactaatctgatttaaattttcagAACG
GAGTCGCCGCCCCATCCGCCACCTCCCAACTCTCCACCCGTGCCCTCGAGGGAATCTTCGAGG
CCACCCACCGTCTCATCTACGGAGCCAAGGACGACTCCGGACAACGTTACCTCGCCTGGTCCGG
ACACTCCGCCCGTGTCGGAGCCGCCCGTGACATGGCCCGTGCCGGAGTCTCCATCCCAGAGAT
CATGCAAGCCGGAGGATGGACCAACGTCAACATCGTCATGAACTACATCCGTAACCTCGACTCCG
AGACCGGAGCCATGGTCCGTCTCCTCGAGGACGGAGACTACCCATATGATGTCCCAGACTACGC
TATGAGCCGTAGACGAAAAGCGAATCCGACAAAACTGAGTGAAAACGCGAAGAAGCTTGCCAAG
GAAGTTGAAAATTAA

HA
EGL-13 NLS

**Smad4_NES::PCKRS(13IPYER)**
ATGGGAATCGACCTCTCCGGACTCACCCTCCAAATGGACAAGAAGCCACTCAACACCCTCATCTC
CGCC**ATC**GGACTCTGGATGTCCCGTACCGGAACCATCCACAAGATCAAGCACCACGAGGTCTCC
CGTTCCAAGATCTACATCGAGATGGCCTGCGGAGACCACCTCGTCGTCAACAACTCCCGTTCCTC
CCGT**CCA**GCCCGTGCCCTCCGT**TAC**CACAAGTACCGTAAGACCTGCAAGCGTTGCCGTGTCTCC
GACGAGGACCTCAACAAGTTCCTCACCAAGGCCAACGAGGACCAAACCTCCGTCAAGGTCAAGG
TCGTCTCC**GAG**CCAACCCGTACCAAGAAGgtaagtttaaacatatatatactaactaaccctgattatttaaattttcagGCC
ATGCCAAAGTCCGTCGCCCGTGCCCCAAAGCCACTCGAGAACACCGAGGCCGCCCAAGCCCAA
CCATCCGGATCCAAGTTCTCCCCAGCCATCCCAGTCTCCACCCAAGAGTCCGTCTCCGTCCCAG
CCTCCGTCTCCACCTCCATCTCCTCCATCTCCACCGGAGCCACCGCCTCCGCCCTCGTCAAGGG
AAACACCAACCCAATCACCTCCATGTCCGCCCCAGTCCAAGCCTCCGCCCCAGCCCTCACCAAG
**CGT**CAAACCGACCGTCTCGAGGTCCTCCTCAACCCAAAGGACGAGATCTCCCTCAACTCCGGAA
AGCCATTCCGTGAGCTCGAGTCCGAGCTCCTCTCCCGTCGTAAGgtaagtttaaacagttcggtactaactaac
catacatatttaaattttcagAAGGACCTCCAACAAATCTACGCCGAGGAGCGTGAGAACTACCTCGGAAAG
CTCGAGCGTGAGATCACCCGTTTCTTCGTCGACCGTGGATTCCTCGAGATCAAGTCCCCAATCCT
CATCCCACTCGAGTACATCGAGCGTTTCGGAATCGACAACGACACCGAGCTCTCCAAGCAAATCT
TCCGTGTCGACAAGAACTTCTGCCTCCGTCCAATGCTCTCCCCAAACCTCTGCAACTACATGCGT
AAGCTCGACCGTGCCCTCCCAGACCCAATCAAGATCTTCGAGATCGGACCATGCTACCGTAAGGA
GTCCGACGGAAAGgtaagtttaaacatgattttactaactaactaatctgatttaaattttcagGAGCACCTCGAGGAGTTCA
CCATGCTCAACTTCTGCCAAATGGGATCCGGATGCACCCGTGAGAACCTCGAGTCCATCATCACC
GACTTCCTCAACCACCTCGGAATCGACTTCAAGATCGTCGGAGACTCCTGCATGGTCTACGGAGA
CACCCTCGACGTCATGCACGGAGACCTCGAGCTCTCCTCCGCCGTCGTCGGACCAATCCCACTC
GACCGTGAGTGGGGAATCGACAAGCCATGGATCGGAGCCGGATTCGGACTCGAGCGTCTCCTC
AAGGTCAAGCACGACTTCAAGAACATCAAGCGTGCCGCCCGTTCCGAGTCCTACTACAACGGAAT
CTCCACCAACCTCTAA

HuSMAD4 NES

**Supplementary Note 2 commercial availability of non-canonical amino acids.**

To the best of our knowledge, at the time of publication three of the photo-caged amino acids used in this study are commercially available. We list some vendors below. We also include existing CAS numbers as well as synonyms for the names of the compounds to help in searching for additional vendors.

**ONBY:**
CAS No. 207727-86-4
O-(2-Nitrobenzyl)-L-tyrosine
(S)-2-Amino-3-(4-((2-nitrobenzyl)oxy)phenyl)propanoic acid

available from:
Fluorochem (product code F447104)
Santa Cruz Biotechnology (product code sc-363295)

**NPY**
CAS No. unknown
6-nitropiperonyl tyrosine
*o*-Nitropiperonyl-*O*-tyrosine
(*S*)-2-amino-3-(4-((6-nitrobenzo[*d*][1,3]dioxol-5-yl)methoxy)phenyl)propanoic acid

Not currently commercially available. Previously available from Sigma-Aldrich (product code 914525)

**MNPY**
CAS No. unknown
Methyl-*o*-nitropiperonyl tyrosine
(2*S*)-2-amino-3-(4-(1-(6-nitrobenzo[*d*][1,3]dioxol-5-yl)ethoxy)phenyl)propanoic acid

Not currently commercially available

**PCK**
CAS No. 1221189-11-2
Methyl-*o*-nitropiperonyllysine
$N^6$-((1-(6-Nitrobenzo[*d*][1,3]dioxol-5-yl)ethoxy)carbonyl)-L-lysine

available from:
Sigma-Aldrich (product no. 915793)

**PCC2**
CAS No.1551078-43-3
S-(Methyl-o-nitropiperonyl)-cysteine
H-L-Cys(MDNPE)-OH
1-[4',5'-(methylenedioxy)-2'-nitrophenyl]ethyl]-L-cysteine

available from:
Iris Biotech GmbH (product code HAA9270)
Chem-Impex (Catalog Number 47699)

**Supplementary Note 3 Fiji script used to analyse activation efficiency of PC-FLP and PC-Cre.**

```
macro "Macro 3 [H]"
{
	title=getTitle();

//SUM PROJECT CHANNELS
	run("Z Project...", "projection=[Sum Slices]");
	selectWindow(title);
	close();
	run("Split Channels");
	selectWindow("C3-SUM_"+title);
```

```
	close();

// CREATE LOOP TO ANALYSE 10 ANIMALS
	for (i = 0; i < 10; i++)
{

//CREATE ROI
	setTool("rectangle");
	run("ROI Manager...");
	waitForUser("Please create a selection");
	roiManager("Add");

//CROP ROI AND CREATE MASK
	selectWindow("C2-SUM_"+title);
	roiManager("Select", 0);
	RoiManager.multiCrop("/home/j/Downloads/stack/", " show");
	selectWindow("CROPPED_ROI Manager");
	run("Duplicate...", "title=mask");
	selectWindow("CROPPED_ROI Manager");
	run("Duplicate...", "title="+i+"_MKATE"+title);
	selectWindow("CROPPED_ROI Manager");
	close();
	selectWindow("mask");
	setAutoThreshold("Li dark no-reset");
	setOption("BlackBackground", true);
	run("Convert to Mask", "method=Li background=Dark calculate black");

	selectWindow("C1-SUM_"+title);
	roiManager("Select", 0);
	RoiManager.multiCrop("/home/j/Downloads/stack/", " show");
	selectWindow("CROPPED_ROI Manager");
	run("Duplicate...", "title="+i+"_CITRINE"+title);
	selectWindow("CROPPED_ROI Manager");
	close();

//APPLY THAT MASK TO THE MKATE IMAGE AND GET HISTOGRAM
	imageCalculator("Multiply create 32-bit",""+i+"_MKATE"+title,"mask");
	selectWindow(i+"_MKATE"+title);
	close();
	imageCalculator("Divide create 32-bit", "Result of "+i+"_MKATE"+title,"mask");
	selectWindow("Result of "+i+"_MKATE"+title);
	close();
	selectWindow("Result of Result of "+i+"_MKATE"+title);
	run("Histogram", "bins=256 use y_max=Auto");
	selectWindow("Result of Result of "+i+"_MKATE"+title);   //asd
	close();

//APPLY THAT MASK TO THE CITRINE IMAGE
	imageCalculator("Multiply create 32-bit",""+i+"_CITRINE"+title,"mask");
	selectWindow(i+"_CITRINE"+title);
	close();
	imageCalculator("Divide create 32-bit", "Result of "+i+"_CITRINE"+title,"mask");
	selectWindow("Result of "+i+"_CITRINE"+title);
	close();
	selectWindow("mask");
	close();
	selectWindow("Result of Result of "+i+"_CITRINE"+title);
	run("Duplicate...", "title=masked");

//THRESHOLD THE MASKED IMAGE AND GET HISTOGRAM
```

```
	selectWindow("masked");
	setThreshold(99000.0000, 1000000000000000000000000000000.0000);
	run("Convert to Mask");
	imageCalculator("Multiply create 32-bit", "Result of Result of "+i+"_CITRINE"+title,"masked");
	selectWindow("Result of Result of "+i+"_CITRINE"+title);
	close();
	imageCalculator("Divide create 32-bit", "Result of Result of Result of 
"+i+"_CITRINE"+title,"masked");
	selectWindow("Result of Result of Result of "+i+"_CITRINE"+title);
	close();
	selectWindow("masked");
	close();
	selectWindow("Result of Result of Result of Result of "+i+"_CITRINE"+title);
	run("Histogram", "bins=256 use y_max=Auto");
	selectWindow("Result of Result of Result of Result of "+i+"_CITRINE"+title);
	close();

//DELETE TEMPORARY STUFF TO RESTART CYCLE
	roiManager("Select", 0);
	roiManager("Deselect");
	roiManager("Delete");
}
}
```

**SUPPLEMENTARY REFERENCES**


1. Chen, Y. Crystal Structure of a Flp Recombinase–Holliday Junction Complex Assembly of an Active Oligomer by Helix Swapping. *Molecular Cell* **6**, 885–897 (2000).
2. Lambert, T. J. FPbase: a community-editable fluorescent protein database. *Nat Methods* **16**, 277–278 (2019).
3. Xi, Z. *et al.* Using a quadruplet codon to expand the genetic code of an animal. *Nucleic Acids Research* **50**, 4801–4812 (2022).
4. Davis, L. *et al.* Precise optical control of gene expression in C elegans using improved genetic code expansion and Cre recombinase. *eLife* **10**, e67075 (2021).
5. Greiss, S. & Chin, J. W. Expanding the Genetic Code of an Animal. *J. Am. Chem. Soc.* **133**, 14196–14199 (2011).
6. O'Shea, J. M. *et al.* Generation of Photocaged Nanobodies for Intracellular Applications in an Animal Using Genetic Code Expansion and Computationally Guided Protein Engineering**. *ChemBioChem* **23**, e202200321 (2022).
